# Formation of clathrin-pits and ATP-independent cholesterol-dependent tubules initiates mechano-regulation on de-adhesion

**DOI:** 10.1101/2023.05.31.543020

**Authors:** Tithi Mandal, Arikta Biswas, Tanmoy Ghosh, Sreekanth Manikandan, Avijit Kundu, Ayan Banerjee, Dhrubaditya Mitra, Bidisha Sinha

## Abstract

Adherent cells ensure membrane homeostasis during de-adhesion by various mechanisms including endocytosis. Although mechano-chemical feedbacks involved in this process have been studied, the step-by-step build-up and resolution of the mechanical changes by endocytosis is not well understood. To investigate this, we study the de-adhesion of HeLa cells using a combination of interference reflection microscopy, optical-trapping and fluorescence experiments. We found that de-adhesion enhanced membrane height fluctuations of the basal membrane in the presence of an intact cortex. A reduction in the tether-force was also noted at the apical side. However, membrane fluctuations reveal phases of an initial drop in effective tension followed by a saturation. The area fractions of early (Rab5-labelled) and recycling (Rab4-labelled) endosomes as well as transferrin-labelled pits close to the basal plasma membrane also transiently increased. On blocking dynamin-dependent scission of endocytic pits, the regulation of fluctuations was not blocked but proceeded uncontrolled. Interestingly, the regulation could not be suppressed by ATP or cholesterol depletion individually but was arrested on depleting both. The data strongly supports pit-formation to be central to the reduction in fluctuations whether in normal or ATP depleted condition. Furthermore, while in normal conditions the contribution of clathrin-mediated endocytosis is clear, under ATP-depleted conditions we propose that cholesterol-dependent pits spontaneously regulate tension.

**Summary:** We show that during de-adhesion, cell edges retract, creating membrane folds and increasing fluctuations. Cells increase the rate of endocytosis to regulate back their membrane fluctuations. This is achieved by forming invaginations. Dynamin-dependent pathways are majorly involved, while cholesterol-dependent ATP-independent mechanisms also contribute.

## Introduction

Sensing and regulating the tension of the plasma membrane is crucial for cells to function effectively, especially when challenged by changes in the local micromechanical environment. Membrane homeostasis ensures the maintenance of membrane tension or conservation of the microscopic fluctuations, providing the plasma membrane’s effective tautness (Morris and Homann, 2001). While enhancing tension can affect endocytosis (Ferguson et al., 2017) and pit-formation (Bucher et al., 2018), as displayed for clathrin-mediated endocytosis (CME), conversely, cells can also alter their endocytosis/exocytosis rates to regulate tension. Studies have addressed the role of membrane trafficking in tension regulation, either in general (Sheetz and Dai, 1996) or focussing on particular pathways such as the CLIC-GEEC (CG) pathway (Thottacherry et al., 2018), CME (Djakbarova et al., 2021) or those using caveolae (Mayor et al., 2014). Flattening of caveolae can regulate tension surge (Sinha et al., 2011) suggesting that pits that participate in endocytosis may directly regulate tension. Since pit-formation and internalization are common to all pathways and sometimes also regulated similarly, it becomes important to understand whether formation of endocytic pits in general can initiate tension regulation or their subsequent internalization is also essential. Addressing such questions would require mechanical perturbation of cells and measuring membrane mechanics and endocytosis state over time. Additionally, perturbing molecules important for pit formation or internalization can further elucidate their individual roles.

To quantify endocytosis, specific cargoes (e.g. transferrin for CME) are usually labelled to quantify its corresponding pathways. However, Rab5 labels early endosomes of a majority of pathways and therefore reports the general state of endocytic trafficking (Bucci et al., 1992). A fraction of endosomes are recycled back to the plasma membrane using a fast-recycling arm, Rab4 (Sönnichsen et al., 2000) which can accumulate if either excess endosomes are formed or the fusion with the plasma membrane is hampered (Jovic et al., 2010). Therefore, following labelled cargo (like Transferrin), as well as Rab5 and Rab4 can collectively provide a description of the trafficking state of the basal plasma membrane.

For perturbing endocytosis, molecules utilized by multiple pathways can be targeted. For example, many constitutive pathways in HeLa cells (e.g. CME, caveolae) require dynamin for the scission of their endocytic pits, making it an important target. Dynamin’s GTPase action can be inhibited by Dynasore, an agent that can prevent scission and thus internalization of pits in these pathways without affecting pit-formation (Macia et al., 2006). However, certain constitutive pathways, like CG, could still be operational even on Dynasore treatment – although its presence in HeLa cell is debated (Thottacherry et al., 2019) while other dynamin-independent constitutive pathways are not well classified. Cholesterol is required very early for the basic clustering of lipids/proteins in several pathways (Klein et al., 1995). The formation of endocytic carriers requires cholesterol for pathways such as CG pathway and caveolae while assembly of clathrin in CME requires adapters like AP2 but is cholesterol-independent (Cocucci et al., 2012; Imelli et al., 2004). Depletion of cholesterol (by MβCD), thus, can arrest the formation of new pits of the CG pathway and caveolae without critically affecting CME. The use of ATP-depletion can reduce both the pinching-off of a majority of pathways as well as the formation of pits in pathways like CME (Schmid and Carter, 1990) and caveolae (Sinha et al., 2011). Thus, ATP depletion could supress the completion of a multitude of constitutive endocytic pathways and therefore could be used to study the role of scission in general. The alterations of these endocytic events need to be checked after imparting mechanical perturbations.

De-adhesion has been often used as a mechanical perturbation in studies on the mechano- chemical feedback in tension regulation (Thottacherry et al., 2018) or the role of blebbing in tension regulation (Norman et al., 2010). Although measurements of apparent tension have been reported during cell spreading (Gauthier et al., 2011; Sinha et al., 2011) similar studies during de-adhesion have not been performed, to the best of our knowledge. Measurement of membrane mechanics can be either done at certain points by optical tweezer based tether pulling or by studying the spontaneous fluctuations of the membrane measured using interference reflection microscopy (IRM). Several studies (Shi et al., 2018; Biswas et al., 2019) have proposed spatial heterogeneities in tension. Multi-point measurements would enable accessing the local distribution of excess membrane undulations or tension, in cells during different phases of de-adhesion .

In this study we mainly employed the IRM-based measurement of effective fluctuation-tension of de-adhering HeLa cells. IRM (Curtis, 1964; Case and Waterman, 2011; Abercrombie and Dunn, 1975; Rädler and Sackmann, 1993; Godwin et al., 1989; Limozin and Sengupta, 2009) primarily provides information about the distance of the basal membrane from the substrate – also called the membrane ’height’. Spatio-temporal measurements of height provides fluctuation amplitude (SDtime), excess area and enables deriving mechanical parameters like fluctuation-tension (Shiba et al., 2016; Biswas et al., 2019).

IRM provides the ability to map fluctuation-tension and get its distribution in the cell and thus differentiate between large global changes and local ones. In this study we have compared changes that are at the level of whole cells (termed cell-wise) and those that show up in the local distribution of tension in single cells. However, while comparing local data, to ensure that the repeated measurements per cell do not falsely strengthen the statistics, we have employed linear mixed models (LMM) as usually used (Herbig et al., 2018; Reichel et al., 2022) to account for the nested grouping of replicate measurements.

Finally, it is important to note certain limitations. Fluctuations reported by IRM are mainly thermal in nature but can have contributions from active (ATP dependent, non-thermal) processes in cells (Turlier and Betz, 2019). A recently developed algorithm (Manikandan et al., 2022, 2020), using a model-free method quantified the effect of active forces on membrane fluctuations (termed activity) and revealed local membrane fluctuations to be weakly active. However, fluctuation-tension solely should not be used for drawing inferences.

We deal with this, primarily, by using direct measurements of fluctuations amplitude. We also compare activity at different time points of de-adhesion to record the level of deviation of the measured fluctuations from equilibrium. Finally, we corroborate the main effect of de-adhesion on fluctuation-tension with apparent membrane tension measurement using the generally accepted optical-trap-based tether extraction method.

Using total internal reflection fluorescence (TIRF) microscopy, we image Rab5 and Rab4 labelled structures, labelling the early endosomes and rapidly recycling endosomes, respectively, to measure how de-adhesion alters endocytosis. We perturb the ability of cells to endocytose to gauge how endocytosis regulates tension and study mechano-regulation on de-adhesion under all such conditions.

## Results

### De-adhesion mediated increase in membrane fluctuations is actin dependent

IRM images reflect the distance of the cellular basal membrane from the glass coverslip and, thus, can be used to study the spatiotemporal basal PM height fluctuations in adherent cells (Biswas et al., 2017). Qualitatively, darker regions were closer to the coverslip while intensity increases with height till ∼ 100 nm and periodically oscillates. Calibration with standards (beads) was performed to quantify the relative heights, followed by selecting regions where the intensity-height conversion was possible – usually regions that are ∼ 100 nm from the coverslip and fall under the first branch of the intensity profile (that displays bands) (Biswas et al., 2017). The selected regions (square membrane patches) were termed first-branch regions (FBRs). Multiple such regions were marked for every cell.

The time series of membrane height fluctuations (obtained from single pixels) were used to get mean height and standard deviation (SD) of height and termed as SDtime. Pixel-wise measurements were averaged over neighbouring pixels to get “FBR-wise” data. SDspace was obtained from a snapshot – comparing height across NxN pixels of any FBR where N was usually 12 in this study (see Methods). These parameters depicted the amplitude of fluctuations. Averaging over all FBRs in a cell and pooling such data from all cells provided the “cell-wise” data. We compared these parameters between control and treated cells to understand how a mechanical perturbation like de-adhesion altered membrane mechanics.

HeLa cells were treated with either a low (0.05% Trypsin-EDTA) or high concentration (0.25% Trypsin-EDTA) of de-adhering solution and imaged by IRM (**Fig. 1a**) to study the changes in membrane fluctuations and mechanics in the basal membrane during de-adhesion. We observed high variability in the time cells took to de-adhere (**Fig. 1b**). To quantify the rate of de-adhesion, the time taken by each cell to reduce their spread area to 67% of the initial was calculated (**Fig. 1c**). We found that, among the various treatments used, Cytochalasin D (Cyto D) – an agent reducing actin polymerization (Schliwa, 1982) of the actin cortex resulted in a very fast de-adhesion. A lower concentration of de-adhering solution of Trypsin (0.05%) was therefore used for Cyto D experiments. This was in line with the understanding that during de-adhesion the contractile cortex caused lateral retraction (Kumar et al., 2019; Sen and Kumar, 2009). At lower trypsin concentration, the process could be slowed down such that the decay times for Control and Cyto D were similar (**Fig. 1c**).

**Figure 1.**
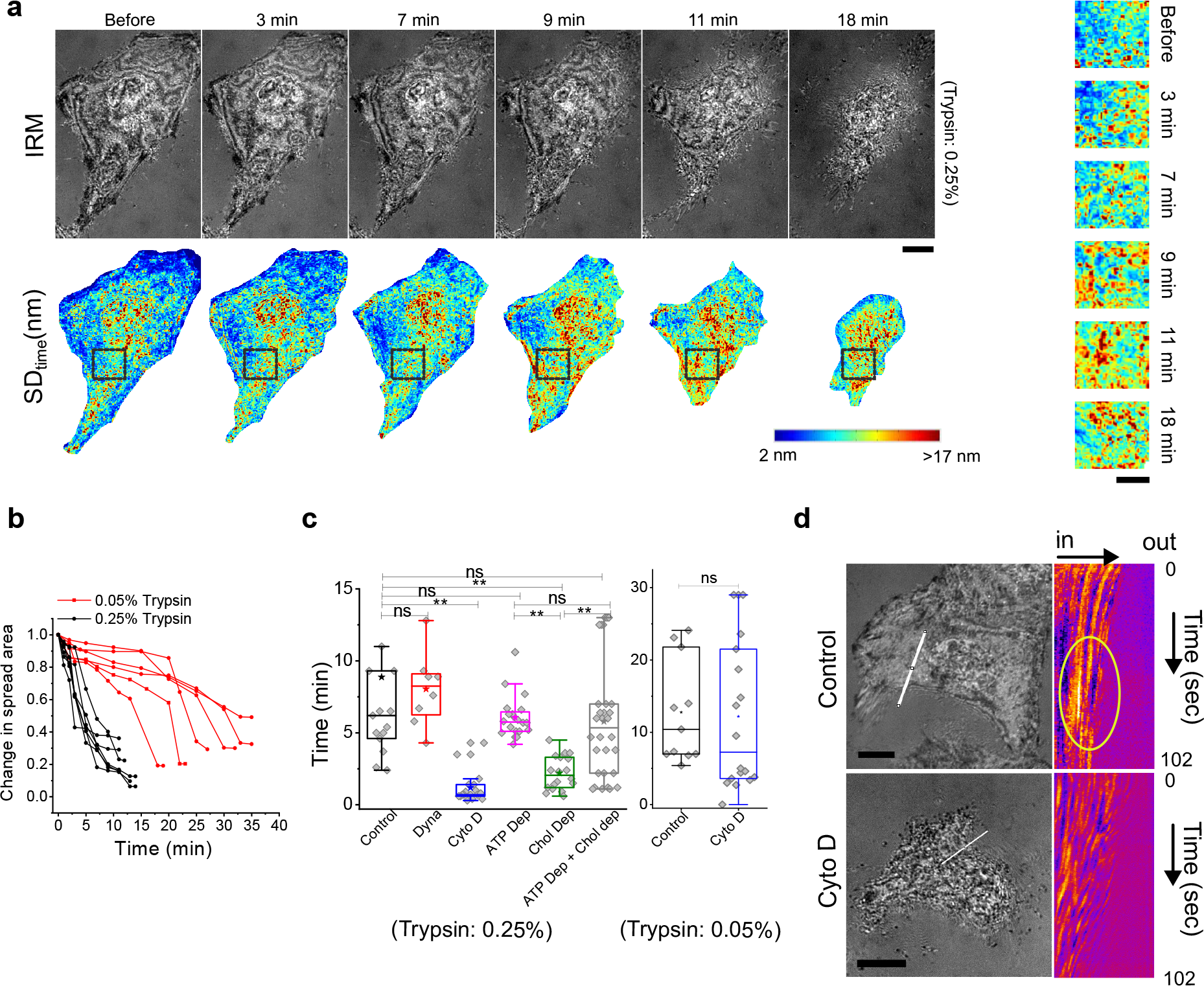
Phases and variability in de-adhesion (a) IRM images of a representative cell before and after Trypsin-EDTA addition at 0.25% (faster) concentrations and corresponding temporal fluctuation map. The time after de-adhering solution addition is mentioned in white at the top right corner. Scale: 10 μm. Zoomed in view of SDtime map (right). **(b)** Profiles of spread area with time after Trypsin-EDTA is added in the two different concentrations. **(c)**Time taken to de-adhere 67% of spread area of cell. **(d)** Representative colour-coded kymographs of IRM intensity of Control and Cyto D treated cell. ROIS drawn perpendicular to the de-adhering front of a cell. Scale: 10 μm

In general, de-adhesion caused an increase in temporal fluctuations (**Fig. 1a** – 7 min), followed by lateral retraction (**Fig. 1a** – 9- 18 min). The increase in fluctuation amplitude, SDtime could be visualized from the maps (**Fig. 1a bottom, Fig. 1a right**). Following the lateral retraction of the edge using a kymograph) of the IRM intensity (along the white line, **Fig. 1d**), we found that as the edges retract inwards, intensity patterns lying inward also moved. Membrane height (IRM intensity) increased in a cluster close to the edge (white oval, **Fig. 1d**) as the retraction progresses. This accumulation faded away with time (section below the oval, **Fig. 1d**). While the membrane undulations were locally enhanced in control cells, such local increase was less prominent in Cyto D treated cells (**Fig. 1d**). Thus, IRM demonstrated that, as de-adhesion progressed, accumulation of membrane height not only increased but could build-up locally due to the cytoskeleton, following which they were also quickly (< min) resolved.

We proceeded to measure the fluctuations and effective membrane-mechanical parameters from the fluctuations during the different phases and over time to quantitatively underpin the effect of de-adhesion on membrane mechanics.

### The initial rise in fluctuations is regulated back

Since the rates of de-adhesion in cells were different, we classified the data in this and following sections based on the level of reduction in spread area. For every cell, we divided the process of de-adhesion into four phases. The “C” phase was defined as the period for which the cell had not been treated with the de-adhering agent (Trypsin). The “P1” phase was demarcated as the slow de-adhesion phase before the exponential fall of the spread area starts. Typically the spread area in this phase remained within 90% of the initial cell spread area. The “P2” phase was described when the spread area exponentially reduced, at least to ∼ 67% of the original value, while the “P3” phase marked out the period when the low spread area had stabilized to ∼ 40-20% of the initial (**Fig. 1b**).

Height fluctuations were measured before (phase: C) and then every few minutes after adding de-adhering medium from image stacks acquired at every time-point, where each movie lasts for ∼ 102 seconds. Only membrane regions that remain adhered through the movies were analysed. On following a representative cell in time (**Fig. 2a**; same cell as shown in **Fig. 1a**) or averaging over a population (**Fig. 2b**), we found that, the amplitude of temporal height fluctuations (SDtime) first increased, followed by a decrease/saturation on de-adhesion. The reverse was observed for fluctuation-tension (**Fig 2a**). Maps of FBR-wise fluctuation-tension (**Fig. 2c**) revealed the lowered tension state of the cell followed by an increase in the regions that remained adhered. Strictly for visualization purposes, we also created a pixel-wise map of tension (**Fig. 2c**). Mapping tension showed a reduction in the initial heterogeneity when cells are at phase P2. The global trend (**Fig. 2b)** was corroborated by the distribution of local (FBR wise) fluctuation amplitude and tension for single cells. Clearly, the changes were greater than the error calculated while averaging the distributions over multiple cells and repeats (**Fig. 2d**).

**Figure 2.**
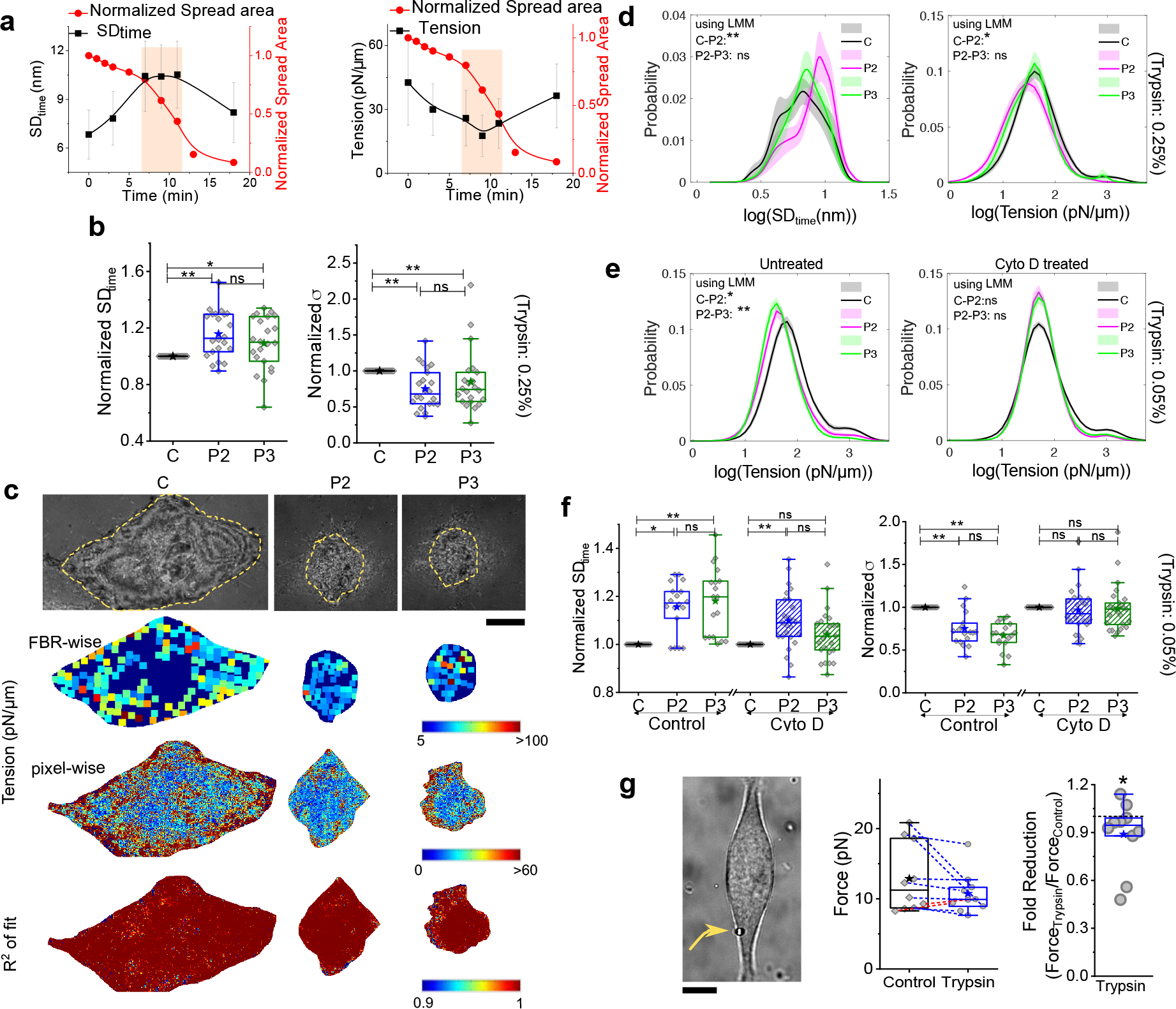
Membrane slacks transiently (a) Representative time series of indicated parameters for a single cell during fast de-adhesion. **(b)** Comparison of normalized amplitudes of fluctuations and tension following the same cells across the indicated phases of de-adhesion involving 22 cells (faster) using FBRs of sizes 4.67 μm^2^(slower) and 0.75 μm^2^(faster). Normalization is performed by dividing any particular cell’s measurement at P2 or P3 by the measurement at C. Table S1 provides a list of the number of FBRs. The size of the FBRs. **(c)** IRM images of a cell at different phases of de-adhesion, and the corresponding FBR-wise tension values mapped back on the cell outline. The dark blue background represents the cell, and coloured boxes denote tension values derived from averaged PSDs from the indicated regions. Lower panel indicates pixel wise tension map and corresponding R^2^ map Comparison of the excess area and activity for faster de-adhesion in single cells. **(d)** Comparison of probability of logarithm of temporal fluctuation and tension across the de-adhesion phases using 0.25% Trypsin-EDTA. **(e)** Comparison of probability of logarithm of tension across the de-adhesion phases of Control (left) and Cytochalasin D (right) treated cells using 0.05% Trypsin-EDTA. **(f)** Comparison of fold change in the mean amplitude of fluctuations and median tension for the same cells followed over time of control and Cyto D treated cells. Normalization is performed by dividing any cell’s measurement at P2 or P3 by the measurement at C. **(g)** Left: Typical image of a cell used for the tether-pulling experiment. The trapped bead is marked out with an arrow. Right: Fold reduction of force after Trypsin addition (higher concentration: 0.25 %). Black * denotes Mann-Whitney U statistical significance test with Bonferroni correction is performed, * denotes p values < 0.016, and ** denotes p value < 0.001. Scale bar =10 μm.

We confirmed that in the absence of de-adhesion media, the fluctuations or tension of these cells did not change over 20 mins (**Fig. S1**). Following the same regions in single cells through de-adhesion also showed the dip and recovery of tension (**Fig. S2**), confirming that regions that remain adhered also went through these changes. Although the spatial height variation (SDspace) and excess area, showed a decreasing trend in contrast to SDtime (**Fig. S3a**), using a gentler substrate detachment reagent – TrypLE (**Fig. S3b**) or at lower trypsin concentration, their trends matched (**Fig. S3 c,d** (**0.05% Tryp**)).

The corresponding entropy-generation rate obtained from fluctuations of de-adhering cells changed mildly(**Fig S3a**) implying that the measured fluctuations did not capture any major enhancement in non-equilibrium activity in the frequencies assayed. Since actin remodelling was expected during de-adhesion, its impact in tension reduction was next studied.

### The initial rise in fluctuations is weaker on cortex disruption

We used Cyto D to weaken the cortical actin. At 0.05% Trypsin spread area reduction rate was similar for Control and Cyto D but cells de-adhered properly. Tension reduction in P2 was not substantial in the presence of Cyto D or reduced filamentous cortical actin (**Fig. 2 e, f**). Although fluctuations were enhanced weakly (than control (**Fig. 2f, S4**) at P2, by P3, unlike in control Cyto D showed similar fluctuations as C. In line with **Fig. 1d** this data also suggests that disruption of cortical actin reduced the accumulation of membrane or enhancement of fluctuations.

De-adhesion only occurs at the basal membrane. To study its effect on the apical membrane, we next extracted membrane tethers from cells and measured tether forces. These were measured on single cells - before and after initiating de-adhesion (**Fig. 2f top, Fig. S5**) within the first 3-7 min. There was a reduction in force for 8/10 cells which showed 2-52% reduction in force (or effectively 4 - 77% reduction in apparent tension).

Next we examned whether the reduction in tension and its subsequent regulation correlated with endocytic activity.

### De-adhesion increases early endocytic and fast recycling endosomes near basal plasma membrane

To quantify endocytosis, we utilized three strategies – labelling the cargo of an endocytic pathway, labelling early endosomes using Rab5 and labelling recycling endosomes with Rab4 (**Fig. 3a**). At first, we followed a cargo of CME – transferrin (Tf) – added in control and de-adhering cells (**Fig. 3b, c**) and found a clear increase in Tf’s incorporation (per μm^2^) in de-adhering cells (**Fig. 3b, c, Fig S6**).

**Figure 3.**
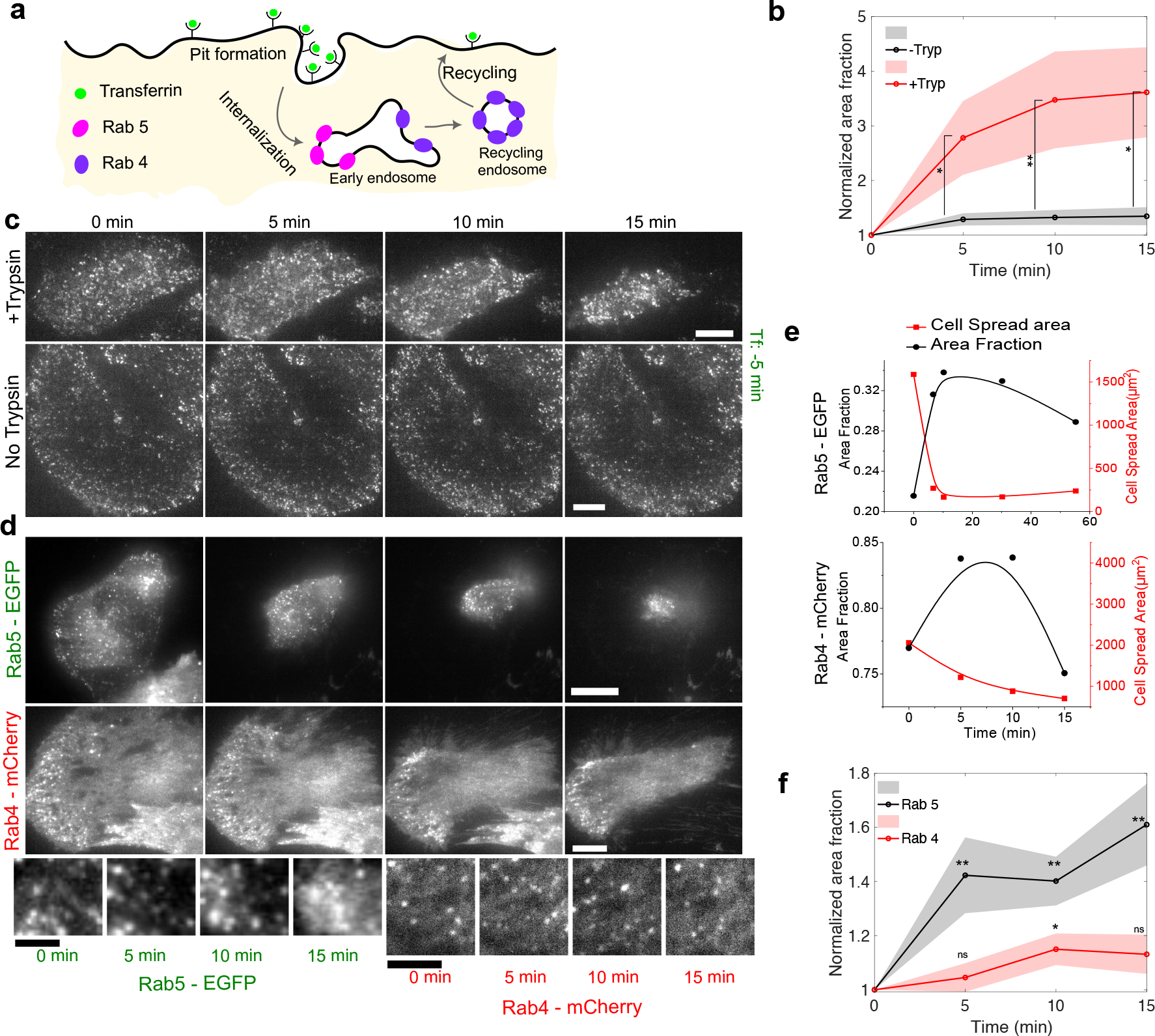
De-adhesion induces endocytosis (a) Schematic of Early endosomes formation and maturation into Recycling endosomes. **(b)** Normalized area fraction of Tf marked puncta followed through time with and without Trypsin. **(c)** Representative TIRF images of HeLa cells followed through time, puncta marked with Transferrin- 568 with 0.25% Trypsin (upper), without Trypsin. Scale bar= 10 µm **(d)** Representative TIRF images of the same HeLa cells transiently expressing EGFP -Rab5 before (0 min) and after administration of de-adhesion media(upper) Scale bar= 10 µm and zoomed in view (Scale bar= 1 µm) and same HeLa cells transiently expressing mCherry -Rab4 before (0 min) and after administration of de-adhesion media(lower)(Scale bar= 10 µm) and zoomed in view (Scale bar= 5 µm). **(e)** Change in area fraction of Rab5 (top) and Rab4 (bottom) as spread area reduces on de-adhesion for a typical single cell. **(f)** Normalized area fraction of Rab5 and Rab5 through different time points before after addition of 0.25 % Trypsin. Table S1 provides the list of the number of cells.

The involvement of the endocytic machinery was next confirmed by labelling early endosomes by Rab5 (Bucci et al., 1992) and the short-loop (fast) recycling endosomes by Rab4 (Sönnichsen et al., 2000). Near the plasma membrane, endosomes mainly contain these two labels (Sönnichsen et al., 2000). The Rab4-containing endosomes emerge from the same endosomes as Rab5 (Sönnichsen et al., 2000) and thus are studied to evaluate the state of endocytic and recycling activity. We imaged EGFP-Rab5 and Rab4 -mCherry using TIRF microscopy (**Fig. 3d, e, Fig S6**). The observed fluorescent puncta were detected as objects and analysed (**Fig S6a)** . Since nearby endosomes could be distinguished from connected ones, we scored for the projected area covered by the endosomes per µm^2^ (termed as area fraction). The area fraction of early endosomes (**Fig. S6**), calculated for whole cells, showed an increase and a subsequent tapering off/decrease (**Fig. 3e**). The increase was anti-correlated initially with the reduction in spread area (**Fig. 3e**). However, it was difficult to discount the effect of heterogenous de-adhesion in the observed trend since some regions had denser features than others. Thus, we looked at regions that stayed on through the observed time scale (15 min). We followed the area fraction of Rab5 for the same sub-cellular region of interest (ROI) (**Fig. 3f**). The rise and saturation were found to be consistent (**Fig. 3f**).

To further validate if endocytosis was ramped up, we analysed clathrin-coated pits (**Fig. S6**). Imaging cells fixed before or after faster de-adhesion. Although an increase and saturation in the area fraction of these pits were observed, the changes were not appreciable.

Together the data showed that triggering de-adhesion reduced tension and cells ramped up their frequency of their endocytosis events. We also know from following fluctuations that the lowering of tension was also stalled or recovered back at later stages of de-adhesion (**Fig. 2b**). To understand if endocytosis affected tension regulation and how, we next used pharmacological agents to block endocytosis and followed treated cells on de-adhesion.

### Blocking dynamin-dependent pathways reduces de-adhesion-triggered endocytosis

We studied the effect of blocking dynamin-dependent pathways by treating HeLa cells with Dynasore (Kirchhausen et al., 2008). Specifically, it does not stop formation of pits (clathrin-coated or caveolae among others) but prevents dynamin’s function in scission of pits (**Fig. 4a**). Instead of the rise in Rab5’s area fraction on de-adhesion, an initial drop was observed with de-adhesion as expected since early endosomes were expected to contain Rab5 (Popova et al., 2013). (**Fig. 4b, c, Fig. S7 a,b**). Failure of fission caused tubes to form, which also contained Rab5 (arrows, **Fig. 4b**, **Fig. S7a**).

**Figure 4.**
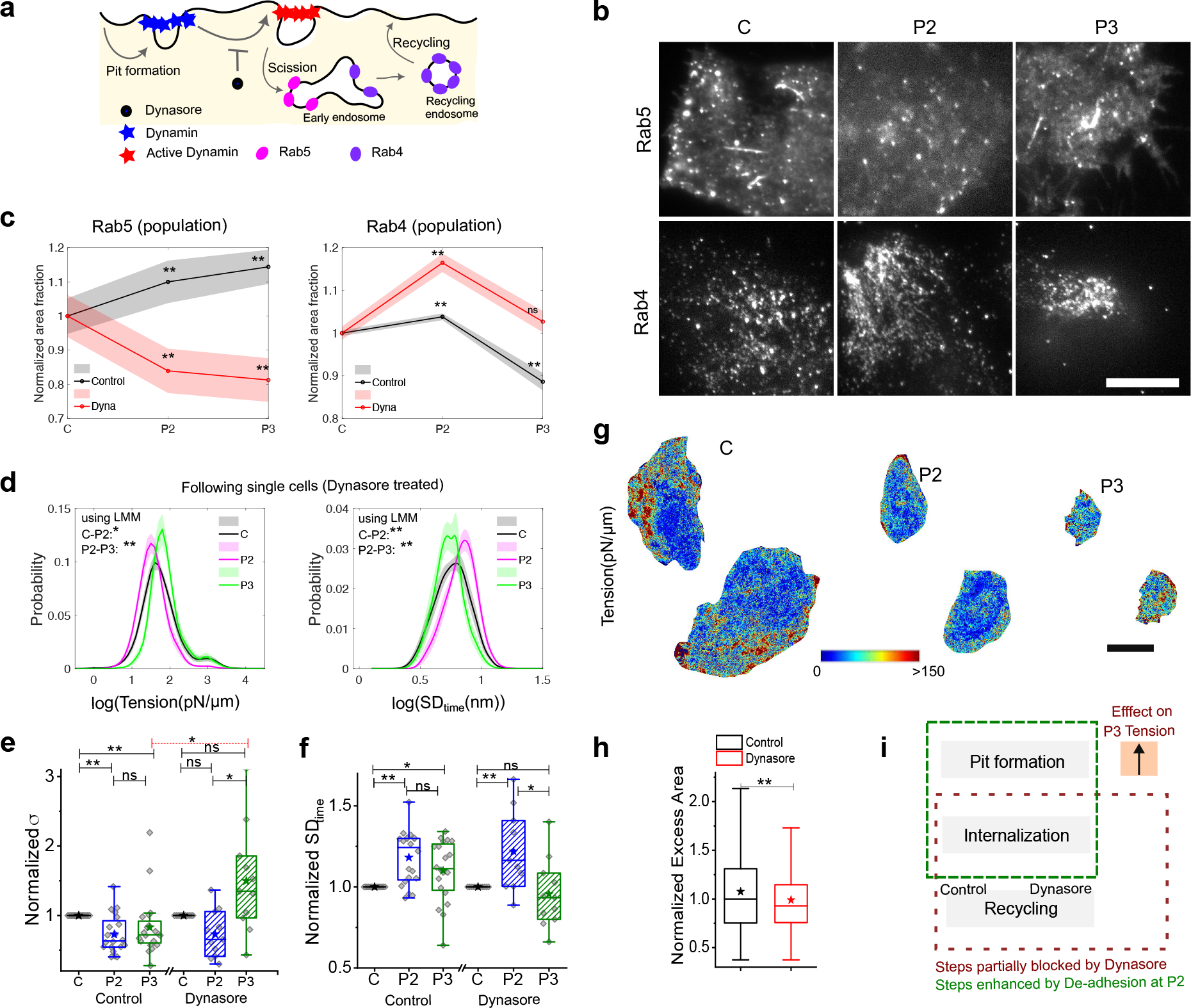
Blocking endocytosis does not stop tension recovery (a) Schematic diagram of pit formation and scission and recycling and inhibition of Dynamin using Dynasore. **(b)** TIRF images of Dynasore treated different cells transfected with Rab5 before and after de- adhesion (top), cells immune-stained with Rab4 before and after de-adhesion (bottom). Scale bar =10 μm **(c)** Normalized area fraction of Rab5 marked early endosomes (left) and Rab4 marked recycling endosomes (right) of control and treated with Dynamin inhibitor- Dynasore. **(d)** Probability distribution of FBR wise log temporal fluctuations and tension of Dynasore treated same HeLa cell followed through different phases of De-adhesion. **(e)(f)** Fold change in cell-averaged parameters comparing each cell with its own measurements at different phases. **(g)** Typical tension map of Dynasore-treated cells in the different phases of de-adhesion. Scale bar =10 μm **(h)** Boxplots of other normalized excess area between Control and Dynasore treated cells. **(i)** Schematic diagram of pit formation, internalisation, and recycling. One-way ANOVA with Bonferroni correction is performed for SDtime since the data is normal. For tension, the Mann Whitney U test is performed and * denotes a p-value < 0.016 (adjusted by group size of 3 per experiment). N= 3 independent experiments. Table S1 provides a list of the number of FBRs.

Rab4 labelling (immunofluorescence) revealed an increase during the de-adhesion (**Fig. 4c, Fig. S7b**). This data suggested that the intermittent accumulation could be due to its inability to fuse normally with the plasma membrane (with reduced tension). This is in line with studies that have reported lowering of recycling on inhibiting Dynamin (Van Dam and Stoorvogel, 2002; Van Dam et al., 2002).

Thus, Dynasore drastically reduced the formation of new early endosomes while also inhibiting the fusion of recycling endosomes with the plasma membrane. Together these indicate Dynasore effectively brought down de-adhesion-triggered endocytosis in the cell. We next studied how such blocking of endocytosis would affect the tension regulation during de-adhesion.

### Blocking dynamin dependent internalization does not stop tension recovery

The single cell distribution of fluctuation amplitude and tension clearly changes as de-adhesion progresses to P2 implying that the tension reduced on de-adhesion of Dynasore treated cells (**Fig. 4d-g, Fig S7d- i**). Interestingly, instead of preventing tension recovery, it was enhanced in Dynasore treated cells (**Fig. 4d**) in comparison to untreated cells (**Fig. 2d, 4e**). Normalized cell averages (**Fig. 4e, f, Fig S7 f**) point to the augmented mechanical regulation operating from P2 to P3 in Dynasore-treated cells. Maps help us visualize this (**Fig. 4g, Fig S7e**). On checking the state of activity (entropy generation rate), we found a lowering of activity on Dynasore treatment (**Fig. S7g**).

To further validate, we performed experiments with a dominant-mutant of Dynamin that was transiently transfected in cells. We found that even in these cells the increase in tension or reduction in SDtime was stronger than in control (**Fig. S8**). Clearly, the tension at P3 was higher than C, unlike in control sets.

The data implies that although endosome formation is reduced, membrane mechanics regulation during de-adhesion is not stopped. This clearly implies formation of pits to have a direct role in the tension recovery. To verify, we next compared the excess areas from IRM images to find the impact of Dynasore on membrane smoothness. There was a significant reduction in the excess area in Dynasore treated cells implying a smoother membrane (**Fig. 4h**) supporting the hypothesis that pit formation can reduce excess area and enhance the effective tension.

Having provided evidence suggesting pit formation as the critical step in endocytosis to cause tension increase, we next aimed to understand the nature of processes used for the initial pit formation. Dynamin-dependent pathways are not limited to CME/caveolae although they are major pathways. To understand the dependency on pathways that use key energy-consuming molecules like dynamin or actin, we next check the regulation’s dependency on ATP. ATP- dependent pit formation are common (CME and caveolae (Sinha et al., 2011) for example) but multiple others like once induced by Shiga toxins are ATP-independent ((Renard et al., 2015).

### ATP-depletion doesn’t block tension regulation

We used ATP depletion prior to de-adhesion to investigate the role of active processes and actin polymerization in the mechanical changes observed during de-adhesion (**Fig. 5a, b**). ATP- depleted cells showed low initial fluctuations but a similar trend of increase in fluctuations followed by a decrease as observed for control cells (**Fig. 5 a, b, Fig. S9a, Table S1**). The effective tension reduced and then increased. Distribution of local values in single cells also captured the changes which were found to be significant. No appreciable change was observed in the area fraction of Rab5 in ATP depleted cells as de-adhesion progressed (**Fig. 5c, d, Fig S9b**).

**Figure 5.**
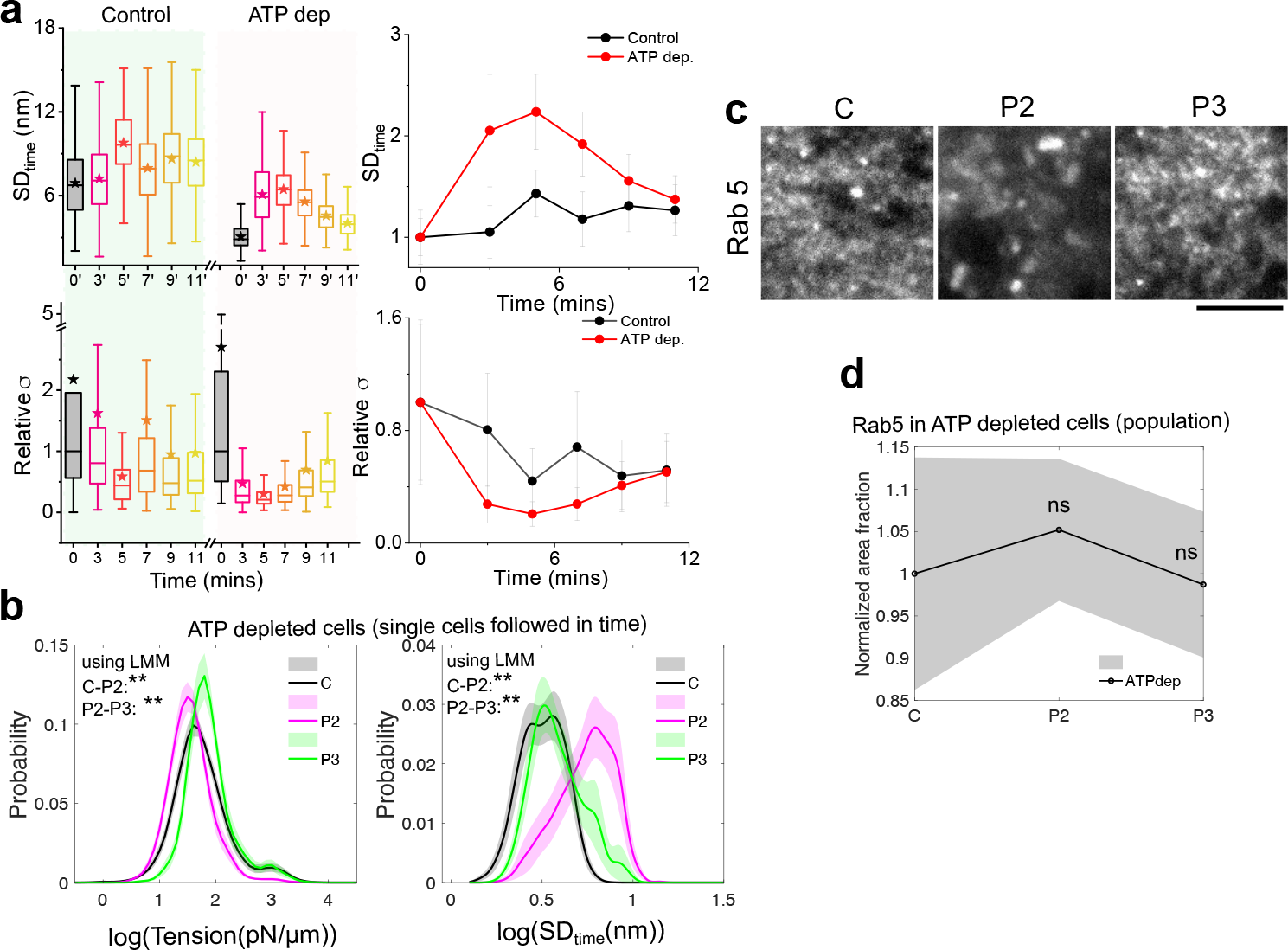
Tension recovery can start without ATP (a) Time series boxplots and median (with MAD as error bar, lower panel) for different parameters for control and ATP-depleted cells on de-adhesion using an FBR size of 4.67 μm^2^. **(b)** Zoomed in TIRF images of different cells transfected with Rab5 before and after de-adhesion triggered in ATP Depleted cells. **(c)** Normalized area fraction of Rab5 in ATP Depleted condition. **(d)** Probability distribution of FBR wise log temporal fluctuations and tension of ATP depleted same HeLa cell followed through different phases of De-adhesion. **(e)** Representative colour-coded kymographs of ATP Depleted HeLa cell ROIS drawn perpendicular to the de-adhering front of a cell.

Therefore, the data here suggests the use of mechanisms that start recovering the tension drop without critically requiring ATP. As the first steps to identify the involved pathway, we next checked in the ATP compromised condition the dependence of tension-regulation on cholesterol. We do so because known pathways (like FEME used to internalize Shiga toxin) using ATP-independent pit formation are cholesterol dependent and depend on the ability of cholesterol to cluster lipids and initiate creation of invaginations (Kovbasnjuk et al., 2001).

### Tension regulation in ATP-depleted cells is cholesterol-dependent

We used ATP depletion and cholesterol depletion as controls (**Fig. 6a, S9e**) to assess the role of cholesterol in tension regulation in ATP-depleted cells during de-adhesion. Comparing single cell distributions or cell-averaged values (**Fig. 6b-d**) showed that while the initial (C-P2) fluctuation (SDtime) increase displayed the same trend as for the controls, fluctuations from P2 to P3 were not reduced efficiently on combined depletion of ATP and cholesterol. Depleting ATP or cholesterol enhanced the recovery of tension (from control) but on dual depletion of ATP and cholesterol there was a further reduction **(Fig. 6a,c, Fig S9 c,d)** (clear from distributions of local measurements and the LMM analysis of the local values (**Fig. 6d**).

**Figure 6.**
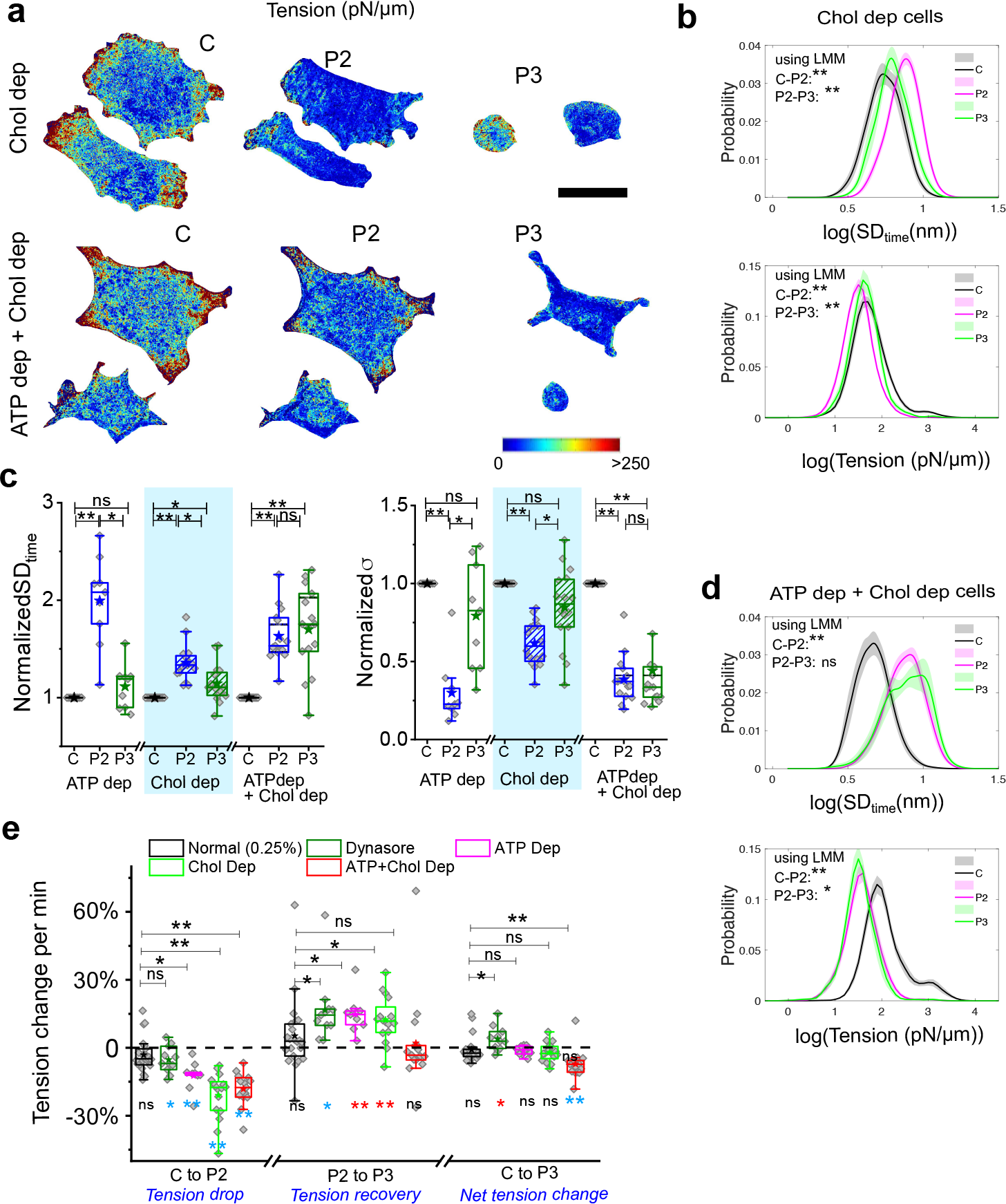
Passive regulation is cholesterol-dependent. (**a)** Representative Tension Map of cholesterol-depleted (Upper Panel) and ATP-depleted as well as cholesterol-depleted cells (Lower Panel) in three phases of de-adhesion. Scale bar = 10 μm. **(b)** Probability distribution of FBR wise log temporal fluctuations and tension of Cholesterol depleted same HeLa cell followed through different phases of De-adhesion. **(c)** Fold change in cell-averaged parameters comparing each cell with its own measurements at different phases. **(d)** Probability distribution of FBR wise log temporal fluctuations and tension of ATP Depleted as well as Cholesterol depleted same HeLa cell followed through different phases of De- adhesion. **(e)**Plot of rate of fractional of tension when cells transit between different phases (mentioned). One-way Anova with Bonferroni correction is performed for SDtime since the data is normal. For tension, the Mann Whitney U test is performed and * denotes p value < 0.016 (adjusted by group size of 3 per experiment). n=3 independent experiments.

Together, we showed (**Fig. 6e**) that the tension drop was significant even on perturbing scission, ATP or cholesterol content and faster in case of the latter two. The recovery was also significant and faster for these perturbations but couldn’t be effective under reduced ATP as well as cholesterol-depleted conditions. Treating tension reached by control cells by P3 phase as a reference, Dynasore treatment effectively tilted the balance towards a larger tension recovery rate while dual depletion of ATP and cholesterol resulted in lower and therefore appreciably slower recovery.

Therefore, the observations strengthen the hypothesis that in the absence of ATP, cells also use cholesterol-dependent processes for mechano-regulation. Since in pathways involved in Shiga-toxin’s endocytosis, toxin-rich tubules have been reported to be formed even in ATP- depleted cells/ giant unilamellar vesicles (Rydell et al., 2014), we next checked if ATP-depleted cells formed tubules when tension was lowered by de-adhesion. For this, we imaged cells transfected with EGFP-CAAX and quantified the membrane cross-section at higher planes in ATP-depleted and ATP and cholesterol-depleted cells using confocal microscopy.

### On de-adhesion, ATP-depleted cells make more cholesterol-dependent tubular invaginations

Cells were transfected with EGFP-CAAX (Madugula and Lu, 2016) to label the plasma membrane and taken through different treatments (**Fig. 7, S10a**). They were subsequently fixed (at 3, 6 min after treatment). Since fixation is not instantaneous, we measured the spread area of cells after fixation to determine the decrease in spread area (**Fig. S10b**) . We observed reduction to ∼ 40% of initial when fixed at 3 min and ∼17% of initial when fixed at 6 min. Therefore, they were classified as P2 and P3. Cells were subsequently imaged in confocal microscopy at a final resolution of ∼ 120 nm in x and y directions (**Fig. 7a, b, S10**). From the intensities obtained from scans under the membrane (**Fig. 7b**) along multiple line regions of interest (ROI), each of length ∼ 4 µm, peaks were detected with larger widths and heights (from basal intensity) than set thresholds from the line scans (**Fig. 7 c**). ATP-depleted cells displayed significantly more internal surface-connected structures than ATP+cholesterol-depleted cells (**Fig. 7d**) at the P2 and P3 phase after de-adhesion. The number of peaks transiently increased with de-adhesion in ATP-depleted cells but, in contrast, decreased for cells also depleted of cholesterol (**Fig. 7d, Fig. S10**). Hence, we concluded that the ATP- depleted cells might gain more cholesterol-dependent tubules on de-adhesion, thereby causing the tension surge.

**Figure 7.**
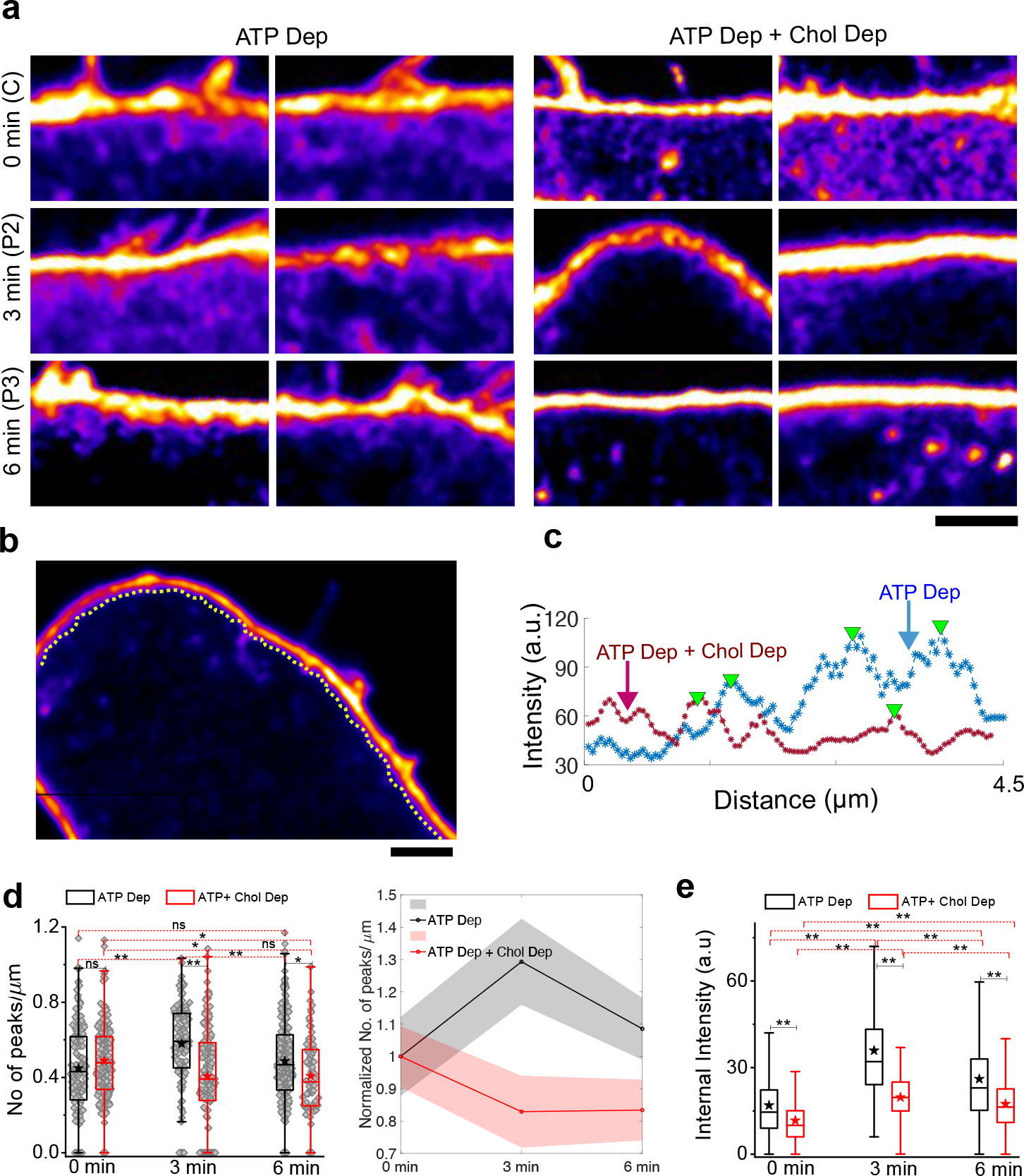
Membrane imaging reveals cholesterol-dependent tubules. (a) Representative zoomed in colour-coded confocal images of cells (ATP-depleted (left) and ATP Depleted as well as cholesterol-depleted (right)). Scale bar = 2 μm. **(b)** Typical line scans are performed parallel to the membrane on the cytosolic side. Scale bar = 2 μm **(c)** Plot profile of typical line scan of ATP depleted and ATP Dep+ Chol Dep with triangles pointing out detected peaks with minimal width and height. **(d)** Box plots (left) and line plot (right) comparing no of peaks in ATP depleted and ATP+ Cholesterol depleted through different time points of de-adhesion. Number of cells= 20,18,28 for 0, 3, 6 min respectively (ATP Depleted), 21,24,21 for 0, 3, 6 min respectively (ATP+cholesterol-depleted). **(e)** Comparison of intensity detected per μm of various 4 μm lines drawn as explained in b. ** denote p value < 0.001 calculated using Mann Whitney U test. Table S1 lists the number of peaks.

In conclusion, we show that de-adhesion induces a tension drop in the whole cell due to an altered adhesion state. Regulation sets in soon and engages endocytic pathways that internalize membrane but also populate a recycling pool. Tension reduction is stalled but must be balanced by recycling back of membrane because perturbations blocking internalization end up enhancing the rate of tension recovery. ATP and cholesterol depletion blocks the tension recovery that blocking internalization or depleting only ATP or cholesterol does not.

## Discussion

In this study, we aimed to understand the basic steps of endocytosis-mediated tension regulation – using de-adhesion as a perturbation and IRM as the primary tool. It should be noted that deriving tension required the use of a model that doesn’t account for the frequency-dependent activity factor. We have evidenced that membrane fluctuations assayed at such small patches (0.75 μm^2^) have overall weak activity (Manikandan et al., 2022) as inferred from measurements of the lower-bound of the entropy generation rate. Using the same algorithm (Manikandan et al., 2022), change in activity is clear when endocytic activity is blocked by Dynasore (**Fig. S7g**) indicating that the technique is capable of resolving small changes. Excess area was observed to be reduced and fluctuation-tension was enhanced (**Fig. S7**). While incorporating the frequency-dependence of activity factor would be ideal while extracting fluctuation-tension, such formulations are currently not yet feasible for easy application to our data.

Hence, direct measurements of fluctuations (SDtime) are first used to comment on the state of the membrane and the effective fluctuation-tension, excess area and functional state of the membrane (endocytic activity) used for further inferences. We also observed that during de-adhesion, the active nature of fluctuations did not significantly increase (**Fig. S3c**). SDtime supported inferences about fluctuation-tension, while apparent tension – measured by optical trapping experiments- validated the initial tension drop on de-adhesion. Furthermore, we not only reported changes in these parameters but also presented functional evidence of the altered physical state. Lowered tension state correlated with enhanced endocytosis (accumulation of Rab5 near the plasma membrane, **Fig. 3f**) and reduced recycling (accumulation of Rab4 near the plasma membrane, **Fig. 3f**). Importantly, we present straightforward evidence of the functional connection between higher fluctuations (and lower effective fluctuation-tension) leading to enhanced endocytosis which in turn leads to reduction of fluctuations. Active models of cell membrane undergoing endocytosis and exocytosis had been used in theoretical work (Rao and Sarasij, 2001) showing that active endo/exocytosis could give rise to an effective tension of the membrane. Their application to IRM data had not been possible but pattern of changes of fluctuations (or effective fluctuation-tension) and Rab5/Tf punctas indicate a similar relationship. Future studies measuring the evolution of fluctuations and endocytosis in same cells would further clarify their local dependence.

Our data first highlights that the local (at the basal membrane) build-up of membrane fluctuations is actin-dependent and not solely dependent on de-adhesion/retraction. The changes in fluctuations were quick to resolve and tension was negligibly altered in Cyto D-treated cells. While this is in line with recent work implicating the cytoskeleton-membrane connections in delaying tension flow and therefore its equilibration, further studies are needed to conclusively prove this.

Identifying the stage when tension recovery initiated was our primary target of investigations. We believe that the data strongly suggests that formation of new invaginations start the tension recovery. Even in the absence of de-adhesion, data in this manuscript (**Fig. S5d**) and reported earlier show that enhanced pit-formation (Dynasore treatment) can lead to reduction of fluctuations and increase in tension (**Fig. S7**). We have also observed that Shiga toxin created tubules and increased tension of ATP-depleted HeLa cells that contained GB3 but did not either create tubules or enhance tension in HeLa cells lacking GB3 (data not shown). The reverse, where flattening of invagination helps cells buffer a tension surge has already been demonstrated. Hence, it is not far-fetched or physically impossible to use invaginations to perform this task. We believe it is possible that clustered lipids/proteins that initiate the pit formation can already start damping the fluctuations although reports suggest so in simulations (Pezeshkian et al., 2017). Pinching-off of endocytic buds or other ATP- dependent mechanisms, therefore, are not critically essential for enhancing tension but required to prevent an excessive rise in tension which could potentially inhibit many membrane processes or start unwanted processes. ATP-dependent machineries, we propose, act as mechano-stats in the cell. They could be vital for keeping tension surges in check rather than being required only for enhancing tension. Our hypothesis is strengthened by the observation (**Fig. 6e**) that the rate of tension-lowering is enhanced in cholesterol-depleted cells, which incidentally, also have a higher fraction of membrane covered by actin-membrane linker – Ezrin (**Fig. S11**).

Finally, some conclusions may be drawn about the relevance of the different pathways of endocytosis in tension regulation during de-adhesion. Assuming the three main pathways to be possibly the CG pathway, CME and caveolae-dependent pathway, cholesterol depletion is expected to block the CG and caveolae-dependent pathways even before pit formation. Cholesterol depletion does not abolish tension recovery or its maintenance close to the control’s response (**Fig. 6e**, C-P3). This shows that CME may significantly contribute mechano- regulation in HeLa cells while passive (ATP independent mechanisms but cholesterol dependent) pathways contribute irrespective of the metabolic state of the cell. We suggest that spontaneously pre-clustered platforms poised for other functions might tubulate on sudden tension reduction and contribute to the passive arm of the regulation.

In conclusion, in this paper, we have presented the effect of de-adhesion on spatiotemporal fluctuations and effective cell membrane mechanics. We have demonstrated that an initial membrane slack is drastically reduced when the actin cortical network is weak. The restoration of the low effective tension involved endocytic pit formation for increasing the tension while recycling for controlling the rise using active as well as cholesterol-dependent passive forms of regulation (**Fig. 8**).

| <u>Resource availability REAGENT</u><br><u>or RESOURCE</u> | SOURCE | IDENTIFIER |
| --- | --- | --- |
| **Pharmacological Reagents** |  |  |
| 2- deoxy-D- glucose | Sigma-Aldrich | Cat# D8375; LOT#<br>WXBB7357V |
| Sodium Azide | Sigma-Aldrich | Cat# 438456; LOT#<br>MKBL3422V |
| Sodium Chloride | Sigma-Aldrich | Cat# S7653; LOT#<br>081M0051V |
| Methyl-β-cyclodextrin | Sigma-Aldrich | Cat# 332615; LOT# STBH0439 |
| Dynasore hydrate | Sigma-Aldrich | Cat# D7693; LOT# 036M4609V |
| ML-141 | Sigma-Aldrich | Cat# SML0407 |
| Cytochalasin D | Sigma-Aldrich | Cat# C2618; LOT# 0000084039 |
| Magnesium Chloride hexahydrate<br>Cryst. Purified | Merck | Cat# 442615; CAS# 7791-18-6; |
| Dimethyl sulfoxide | Sigma-Aldrich | Cat# 276855; LOT# SHBC3339V; |
| HEPES | Sigma-Aldrich | Cat# H3375; LOT# SLBN0452V |
| Gelatin | Sigma-Aldrich | Cat# G2500; LOT# SLBX2973 |
| Glycine | Merck | Cat# 56-40-6 |
| TritonX-100 | Sigma-Aldrich | Cat# 101371902 |
| Recombinant Anti-Rab4 antibody<br>[ EPR3043] | Abcam | Cat#ab109009<br>RRID# AB_10887396 |
| Goat Anti -Rabbit IgG H&L, Alexa<br>Fluor 488 | Abcam | Cat#ab150077<br>RRID# AB_2630356 |
| **Cell culture** |  |  |
| Cell line (HeLa) | ATCC | Cat# CCL-2; RRID# CVCL_0030 |
| Dulbecco's Modified Eagle's Medium (DMEM, High Glucose) | Gibco | Cat# 11965092 |
| Fetal Bovine Serum, certified, heat-inactivated, US | Gibco | Cat# 10082147 |
| Antibiotic-Antimycotic (100X) | Gibco | Cat# 15240062 |
| TrypLE Express | Gibco | Cat# 12605-028; LOT# 2323753 |
| Trypsin-EDTA (0.05%), phenol red | Gibco | Cat# 25300062; LOT# 2193180 |
| Trypsin-EDTA (0.25%), phenol red | Gibco | Cat# 25200072 |
| Paraformaldehyde | Sigma-Aldrich | Cat# P6148 |
| Phosphate Buffered Saline | Sigma-Aldrich | Cat# P3813 |
| Dulbecco's Modified Eagle's Medium (DMEM, Phenol Red free) | Gibco | Cat# 21063-029; LOT# 2239707 |
| Opti-MEM (Reduced serum medium) | Gibco | Cat# 31985-070; LOT# 2192861 |
| Lipofectamine 3000 Transfection kit | Invitrogen | Cat# L3000-015; LOT# 2145954 |
| **Plasmids** |  |  |
| mCherry- Clathrin LC-15 | Addgene | Plasmid# 55019 |
| EGFP-Rab5 | Addgene | Plasmid# 49888 |
| EGFP-CAAX | Addgene | Plasmid# 86056 |
| **For calibration** |  |  |
| NIST Traceable Particle Size Standard, 60 µm | Bangs laboratories | Cat# L130806L; LOT# 11247 |
| **Software and algorithms** |  |  |
| ImageJ (FIJI) | NIH | N/A |
| Origin | OriginLab Corporation | N/A |
| MATLAB | The Mathworks, Inc. | N/A |
| Labview | National Instruments |  |

**Figure 8.**
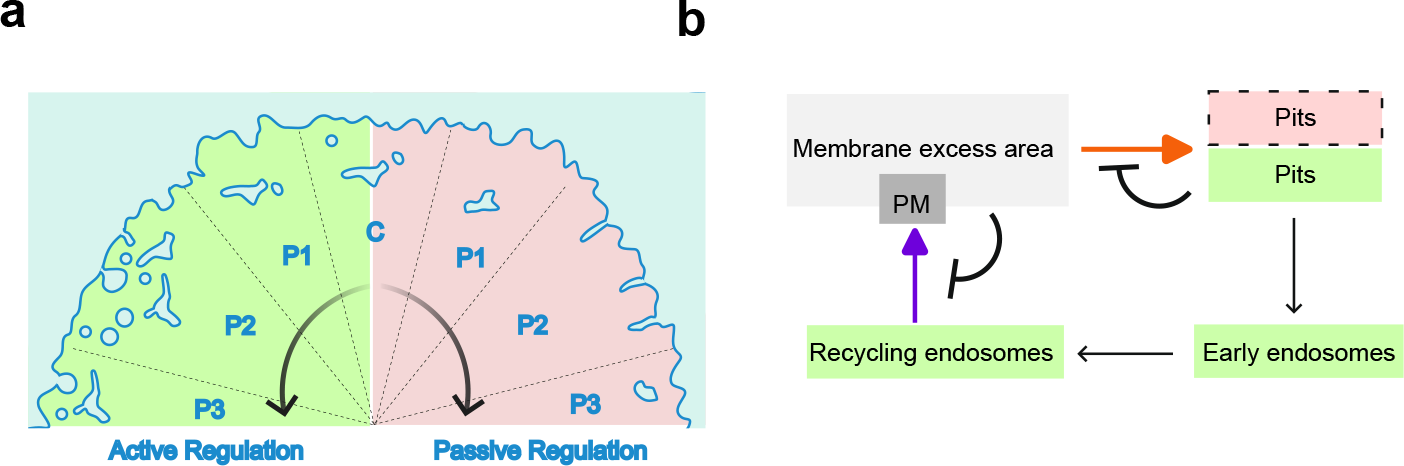
Schematic Diagram. (a) Membrane remodelling by active and passive regulation. Schematic shows that membrane fluctuations enhance in the P1 and P2 phase. However, while in passive condition (low ATP), cholesterol-dependent tubules reduce the membrane fluctuations, in normal conditions, active regulation entails formation of pits and their internalization in the P2 phase which transiently accumulate in early and recycling endosomal structures till the tension enhances back and fusion of recycling membrane keeps the increasing tension in check **(b)** Regulation of plasma membrane excess area by formation of pits, early and recycling endosomes. Schematic depicts that higher membrane excess area at the PM favours formation of pits as observed in this study. Such pits can be static or actively internalized to add to the the early endosomal pool as shown in this study. Part of the early endosomal pool get converted to the recycling pool which is depleted when some structures fuse back with the PM. At higher membrane excess area, this fusion is disfavoured which can cause accumulation of the recycling endosome as shown in a and observed in this study.

## Lead contact

More detailed information and request for the resources should be directed to and will be fulfilled by the lead contact, Bidisha Sinha.

## Material availability

New materials and methods used in these studies will be available upon request to Bidisha Sinha.

## Data and code availability

Data and codes used in this study for analysis purposes will be available upon request to the lead contact, Bidisha Sinha.

## Methods

### Cell line

HeLa cell line (CCL-2, ATCC) was used to perform all the experimental studies.

### Cell culture

HeLa cells were grown in Dulbecco’s Modified Essential Medium (DMEM, Gibco, Life Technologies, USA) with 10% foetal bovine serum (FBS, Gibco) and 1% Anti-Anti (Gibco) at 95% humidity, 5% CO2 and 37 °C. Experiments were always performed after 16-18 h of cell seeding.

### Pharmacological treatments

To de-adhere cells from the substrate, HeLa cells were incubated with 0.05% or 0.25% Trypsin-EDTA solution (Gibco) at 37 °C on the onstage microscope incubator. To inhibit all dynamin-dependent endocytic pathways, we incubated cells with Dynasore hydrate (80 μM; Sigma) in serum-free media for 20 min (Kirchhausen et al., 2008; Barrias et al.; Macia et al., 2006). HeLa cells are incubated with ML-141 (10 μM; Sigma) for 30 min in serum-free media to inhibit CLIC-GEEC endocytic pathways (Thottacherry et al., 2018). For ATP depletion, cells are incubated for 1 h with 10 mM Sodium Azide (Sigma-Aldrich) and 10 mM 2- deoxy-D-glucose (Sigma-Aldrich) dissolved in M1 media composed of 150 mM NaCl (Sigma-Aldrich), 1 mM MgCl2 (Merck), 20 mM HEPES (Sigma) (Zha et al., 1998; Biswas et al., 2017). 10 mM Methyl-ß-cyclodextrin (Sigma-Aldrich) is used in serum-free media for 50 min to deplete cholesterol (Biswas et al., 2019). For inhibiting Actin filament polymerization, cells were kept in 5 μM Cytochalasin D (Sigma-Aldrich) for 1 h in serum-free media (Biswas et al., 2017). Cells are de-adhered by TrypLE Express (Gibco). This is used as an alternative to Trypsin-EDTA for de-adhesion experiments (Thottacherry et al., 2018). All the treatments are incubated at 37 °C temperature inside the incubator. During imaging and de-adhesion all tretaments were maintained at the same concentration.

### Immunostaining

For immunostaining, cells were first fixed with 4% paraformaldehyde (Sigma-Aldrich) for 15 min and subsequently washed twice with phosphate-buffered saline (PBS, Sigma-Aldrich). Next, cells were incubated in 0.1 M glycine (Sigma-Aldrich) for 5 min and then washed again with PBS. Triton-X was used for 2 min and then washed with PBS. For blocking, cells were incubated with 3 ml of 0.2% Gelatin (Sigma-Aldrich) solution for 3 h at room temperature. Primary antibody treatment was done with Recombinant Anti-Rab4 antibody (Abcam) at 1:200 dilution in Gelatin and kept overnight at 4°C to mark recycling endosomes. Goat Anti - Rabbit IgG H&L, Alexa Fluor 488 secondary antibody (Abcam) was used at 1:500 dilution at Gelatin for 2 h after washing with PBS. Subsequently, cells were imaged in 2 ml of PBS.

### Transfection

EGFP-Rab5 (Addgene) was a gift from Arnab Gupta. Cells were transfected with 0.5 μg of Rab5 plasmid DNA to label early endosomes respectively by using Lipofectamine 3000 (Invitrogen). EGFP-CAAX (Madugula and Lu, 2016) was a gift from Lei Lu. Cells were transfected with EGFP-CAAX (Addgene) plasmid to mark the membrane. Cells were transfected with 1 μg of mCherry-Clathrin LC-15 (Addgene) to label clathrin-coated vesicles. Other treatments, if required, were performed 16 h after transfection.

Fixation of cells was performed by using 4% paraformaldehyde (Sigma-Aldrich) for 15 min at 37° C temperature.

### IRM Imaging

Cells were imaged in Nikon Eclipse Ti-E motorized inverted microscope (Nikon, Japan) equipped with adjustable field and aperture diaphragms, 60X Plan Apo (NA 1.22, water immersion) and a 1.5X external magnification on an onstage 37 °C incubator (Tokai Hit, Japan). Either an EMCCD (Evolve 512 Delta, Photometrics, USA) or an s-CMOS camera (ORCA Flash 4.0, Hamamatsu, Japan) was used for imaging. A 100 W mercury arc lamp, an interference filter (546 ± 12 nm) and a 50-50 beam splitter were used (Biswas et al., 2017). For IRM, movies consisted of 2048 frames (19.91 frames/s, EMCCD and 20 frames/s for s- CMOS) recorded at EM 30 and an exposure time of 50 ms.

### Optical trap experiment

For optical trap-based tether-pulling experiments, a 2 µm polystyrene bead was trapped by focusing a 1064 nm Laser (Coherent, Sweden) of 1000 W power at source using a 100x objective. The back-aperture of the objective was over filled after beam expansion and using mirrors and a 50/50 beam-splitter for beam manipulation. The trap stiffness (k) was calibrated from trajectories of the trapped bead using the equipartition approach 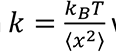 where x is the displacement of the bead from the trap centre, kB is the Boltzmann constant and T the temperature in the Kelvin scale. For analysis, the bead was detected as an object (MATLAB, Fiji), and its centre was tracked with time. The bead’s displacement from the centre of the trap and the spring constant of the trap was used to get the force (*F* = −*kx*). For every cell, a tether was pulled at a constant velocity of 0.5 µm/s up to a distance of 40 µm (LabVIEW, National Instruments, USA). Tether force was calculated from the average bead position during the period when it was parked with the tether pulled (**Fig. S3c**) for ∼ 50 sec. Imaging was done at 200 frames per sec. The apparent membrane tension of the apical section of the cell was derived from the force using the Canham-Helfrich equation 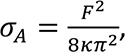, where κ is the bending rigidity and taken to be 15 kBT (Cuvelier et al., 2005), and σA denotes the apparent membrane tension.

### TIRF Imaging

For TIRF Microscopy, an inverted microscope (Olympus IX-83, Olympus, Japan) was used with a 100X 1.49 NA oil immersion TIRF objective (PlanApo, Olympus). An s-CMOS camera (ORCA Flash 4.0, Hamamatsu, Japan) and 488 nm, as well as 561 nm laser sources, were used. Images were acquired using an exposure time of 300 ms with ∼70 nm penetration depth.

### Confocal Imaging

For confocal imaging Leica confocal microscope (Leica SP8) was used with a 63X oil objective lens (NA 1.4). A step size (in z) of 250 nm is used for imaging with a pixel size of 45 nm and deconvoluted (Leica Lightning Software).

### Analysis of IRM images

The intensity of IRM images was converted to height (wherever applicable) as reported (Biswas et al., 2017). The amplitude of spatial undulations spatial (SDspace) was obtained from the standard deviation (SD) of relative heights across all pixels in an FBR after averaging it over 20 frames. For obtaining the amplitude of temporal fluctuations, SDtime, the SD of the relative heights over 2048 frames in each pixel was calculated and averaged across all pixels in an FBR. The power spectral density (PSD) of individual pixels was obtained from the temporal relative height time series using either the FFT method or the covariance method (MATLAB). PSDs of all pixels in an FBR were averaged to obtain the PSD for that FBR. For obtaining mechanical parameters, the PSDs were fitted to *PSD*(*f*) =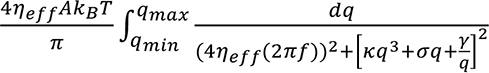 (Helfrich, 1973; Alert et al., 2015; Biswas et al., 2017) where active temperature (A), effective cytoplasmic viscosity (ριeff), confinement (γ) and membrane tension (σ) were used as fitting parameters. The bending rigidity (κ) was fixed at 15 kBT (Simunovic and Voth, 2015). For obtaining maps, PSDs of every pixel were fitted to the theoretical model. For obtaining excess area fraction 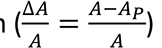 over an FBR, the flat area of the FBR is taken as AP (= L^2^ when patch/FBR is a square of side L) and *A* is the sum of all *dA* calculated for each pixel by comparing the height at that pixel with its neighbours (*dA* = 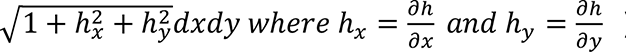 (Deserno, 2007). The excess area is represented in figures as the percentage excess area 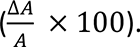.

The activity per FBR is calculated as the lower bound of the entropy generation rate at each FBR. For obtaining the entropy generation rate, data pooled from all pixels of the FBR were taken through dimensional reduction using principal component analysis. The time series (2048 frames) of one of the principal components were built and analyzed as described recently (Manikandan et al., 2022) using the short-time inference scheme reported earlier (Manikandan et al., 2020).

### Fluorescence Image analysis

For endosomal counting (MATLAB), first, a Gaussian blur operation was performed to spatially average out the image. The gaussian-blur is next subtracted from the original image, thus enhancing local contrast. The subtracted image is normalized between 0 to 1, and thresholding was performed using the appropriate threshold, resulting in a binary image. Next, a mask was applied over the image to select the cell. Single pixels were removed using serial erosion and dilation, and subsequently the binary image was used to detect objects. The area fraction is calculated by dividing the total area of early endosomes by the total area of the ROI or cell.

### Counting tubules from confocal z-slices

For counting tubules, Z-stack images of cells were captured. Line-scans were drawn at the cortex regions of cells (ImageJ/Fiji), and the intensities of these line scans were taken from the intensity plot profile of each line-scan to plot the internal intensity. Peak analysis (MATLAB) involved evaluating the intensity line scans for peaks of minimum prominence of ∼ 6 and width of ∼ 5.

### Statistical Analysis

Every IRM experiment was preceded by imaging beads. Every experiment was repeated at least thrice involving multiple cells and FBRs (**Table S1**). A Mann-Whitney U test was performed for statistical significance testing (ns denotes p > 0.05, * denotes p < 0.05, ** denoted p < 0.001). When indicated, a linear mixed effect model (LMM, MATLAB) was also used (using the fixed effect of time and random effects grouped under replicate set number of the experiments and cell number) to quantify the significance of the observed changes in logarithm of tension values (**Table S2**). This helped avoid the effect of the high sample size of FBRs that could influence hypothesis testing. LMM has been used for comparisons in other high-sampling mechanical measurements (Reichel et al., 2022; Herbig et al., 2018).

## Acknowledgement

BS acknowledges support from Wellcome Trust/DBT India Alliance fellowship (grant number IA/I/13/1/500885), SERB (grant number SERB_CRG_2458) and CEFIPRA (grant number 6303- 1). The authors are grateful to CSIR and IISER Kolkata for providing scholarships to TM and AB, respectively. TG thanks CEFIPRA for providing his fellowship. The authors are also thankful to the DBT Wellcome Imaging Facility (IA/I/16/1/502369) for confocal imaging and the DIRAC supercomputing facility for tension mapping.

## Author Contribution

Tithi Mandal: investigation, formal analysis, data curation, writing (original draft), Arikta Biswas: investigation, methodology, software, formal analysis, writing (original draft), Tanmoy Ghosh: methodology, software, investigation, formal analysis, Sreekanth Manikandan: resources, Avijit Kundu: methodology, Ayan Banerjee: methodology, resources, Dhrubaditya Mitra: resources, Bidisha Sinha: conceptualization, methodology, writing (original draft), funding acquisition. All authors edited the manuscript.

