## Supplemental Material for "Formation of clathrin-pits and ATP-independent cholesterol-dependent tubules initiates mechano-regulation on de-adhesion"

**LIST OF FIGURES**

**Figure S1**. Fluctuations and tension of adhered cell - over time

**Figure S2.** Following the same regions through de-adhesion

**Figure S3.** Fluctuations during de-adhesion with different strengths of de-adhering agent.

**Figure S4.** Fluorescence analysis, Transferrin, Early and Late Endosome imaging, Clathrin puncta analysis

**Figure S5.** Early and Late Endosome and Membrane Parameters on blocking endocytosis

**Figure S6.** Membrane fluctuation, Tension, and tension maps on Cyto D treatment

**Figure S7.** Membrane fluctuation, Rab 5 area in ATP Depletion, Parameters and tension maps on ATP and cholesterol depletion and both ATP – Cholesterol depletion.

**Figure S6**. Membrane images marked with EGFP-CAAX

**Figure S9.** Enhanced surface ezrin on cholesterol depletion

**Figure S10.** Fluctuations and Tension of Dynamin2(K44A)-mCherry transfected cell

**Table S1.** List of statistical parameters for all figures

**Table S2.** Values obtained from LMM analysis of FBR-wise comparisons

**Extended Methods :** Details about Linear mixed models.

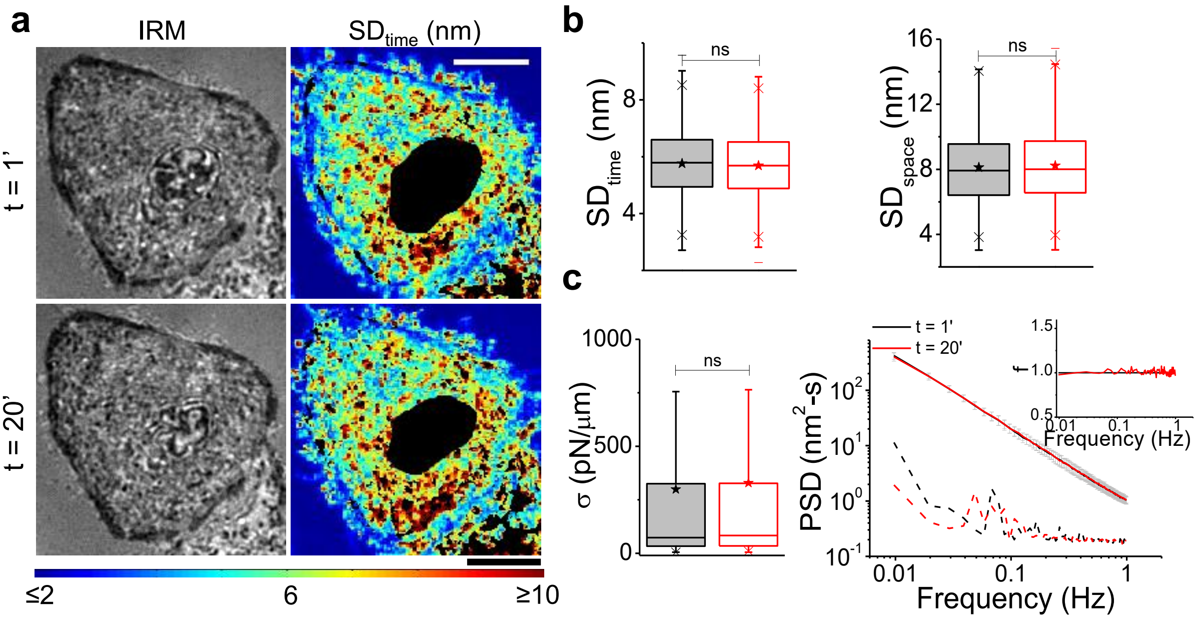

***Figure S1*. *Fluctuations and tension of adhered cell - over time.* (a)** *Representative IRM and* $SD_{time}$ *map of adhered HeLa cell imaged at 1 and 20 min. No de-adhesion media was administered.* ***(b)*** *Temporal and Spatial fluctuations.*

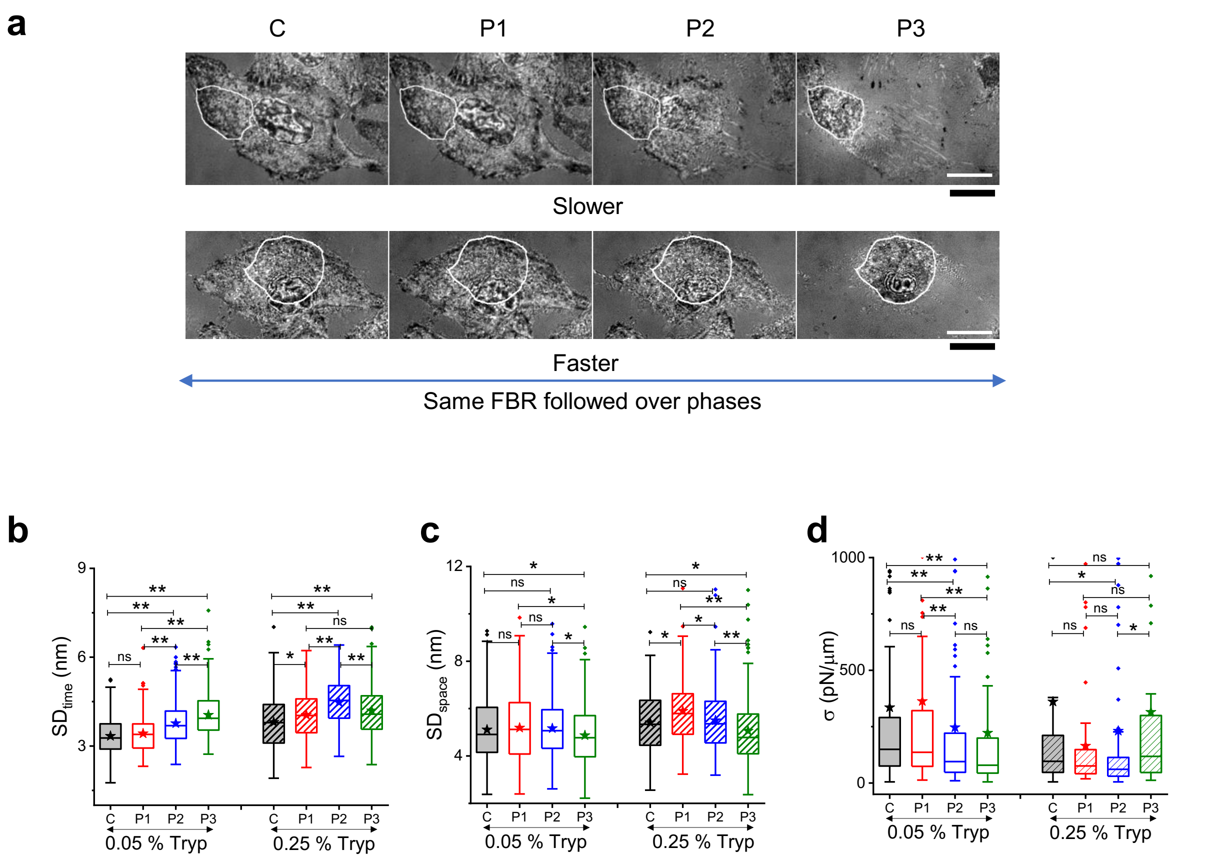

***Figure S2. Following the same regions through de-adhesion. (a)*** *Representative IRM images of HeLa cells during de-adhesion. White outlines mark out the region followed over time.* ***(b)*** *Comparisons of temporal fluctuations and* ***(c)*** *spatial undulations.* ***(d)*** *tension for cells at different phases and for slower/faster de-adhesion.*

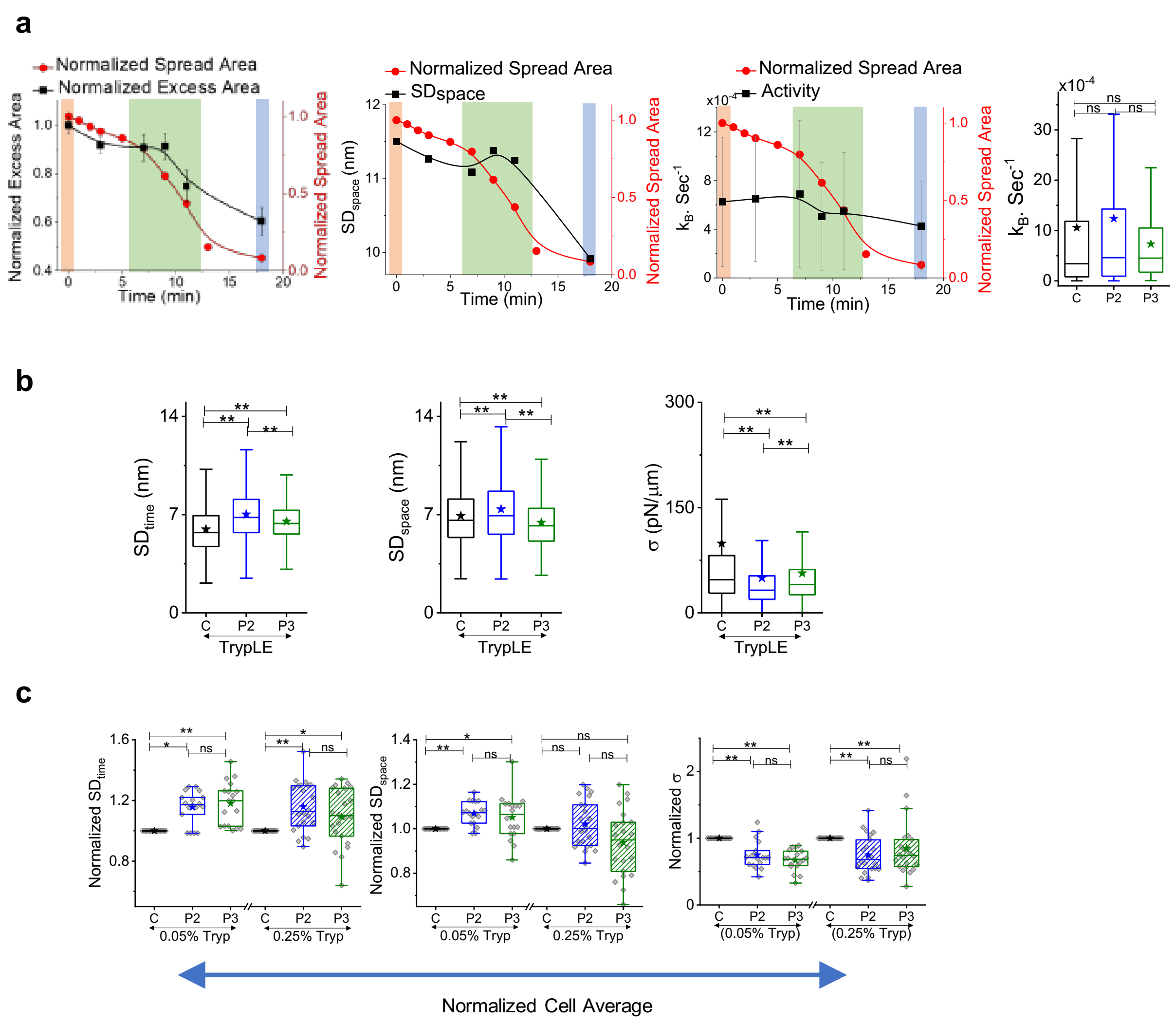

***Figure S3. Fluctuations during de-adhesion with different strengths of de-adhering agent. (a)*** *Representative time series of indicated parameters for a single cell during fast de-adhesion(left) Box Plot of Activity in different phases of de-adhesion (Right).* ***(b)*** *On using milder de-adhesion reagent TrypLE.* ***(c)*** *Cell-wise comparison of fluctuations and tension..*

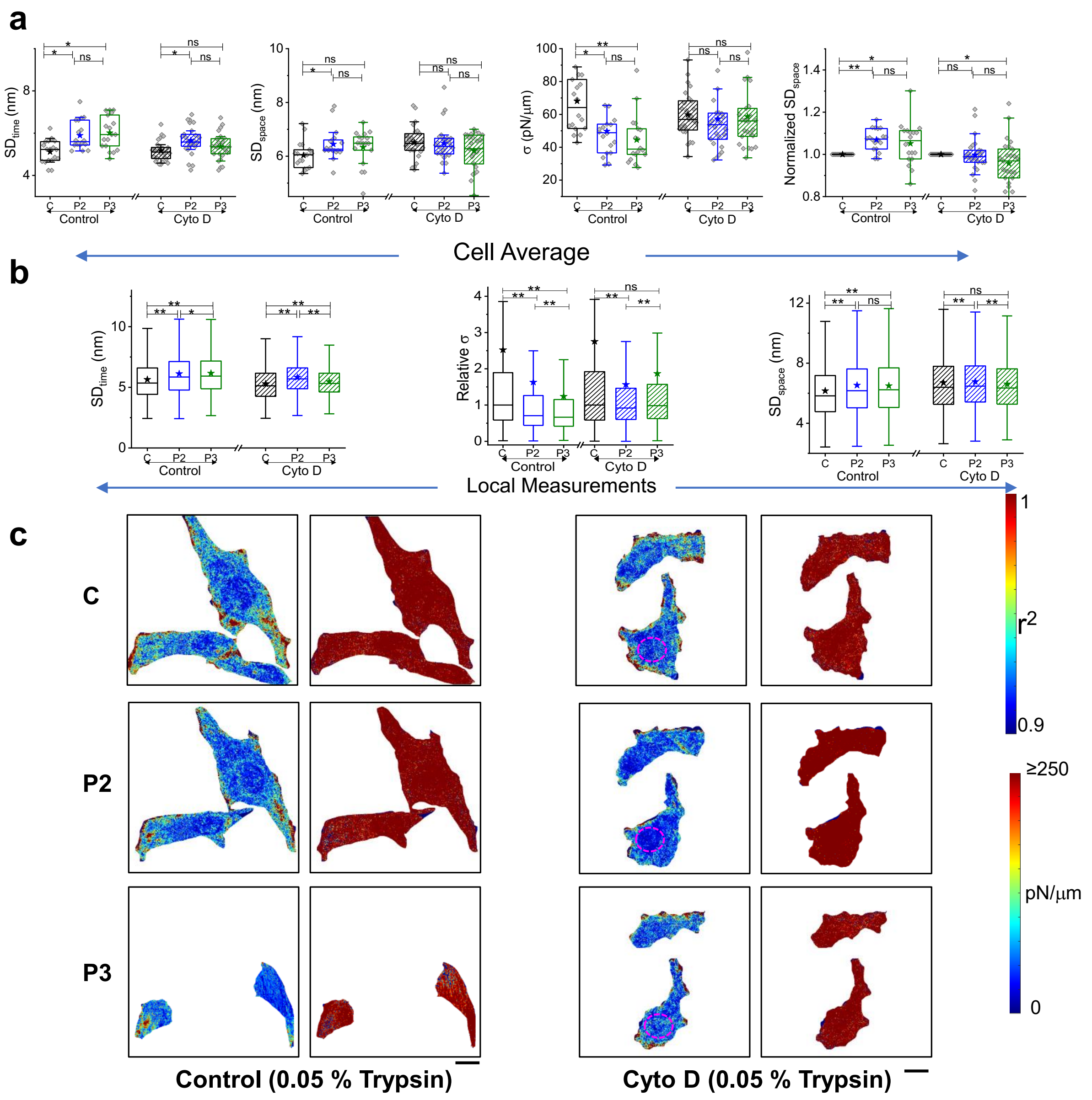

***Figure S4. Membrane fluctuation, Tension and tension maps on Cyto D treatment. (a)** Cell-wise comparison of fluctuations and tension. **(b)** Local measurements of temporal fluctuation, relative tension, spatial undulation at different phases of de-adhesion in control and Cyto D treated cells **(c)** Typical tension maps of control and Cyto D treated conditions at different phases of deadhesion. Scale bar represents 10 μm.*

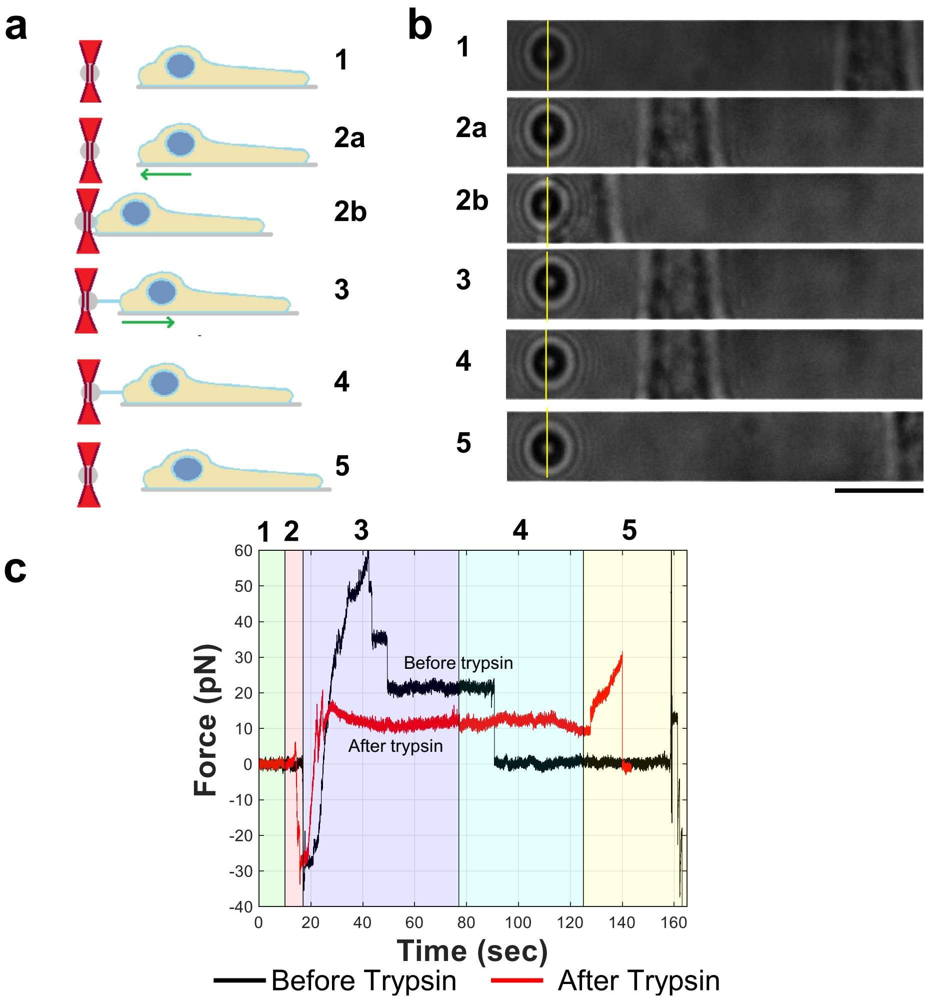

***Figure S5. Optical trapping. (a)*** *Left: Schematic depicts different phases of the optical trapping experiment 1: measuring bead fluctuations away from cell 2a: cell approaching the bead; 2b: Cell and bead interacting; 3: cell pulled away; 4: imaging while bead is parked; 5: moving bead away to rupture the tether to ensure presence of single tether.* ***(b)*** *the brightfield images of the different phases have been put; (**c**) Force vs. time plot shows the force evolution in the different phases. Note that for the black curve, tether breaks while in the waiting phase and for the red curve in breaks on being pulled again.*

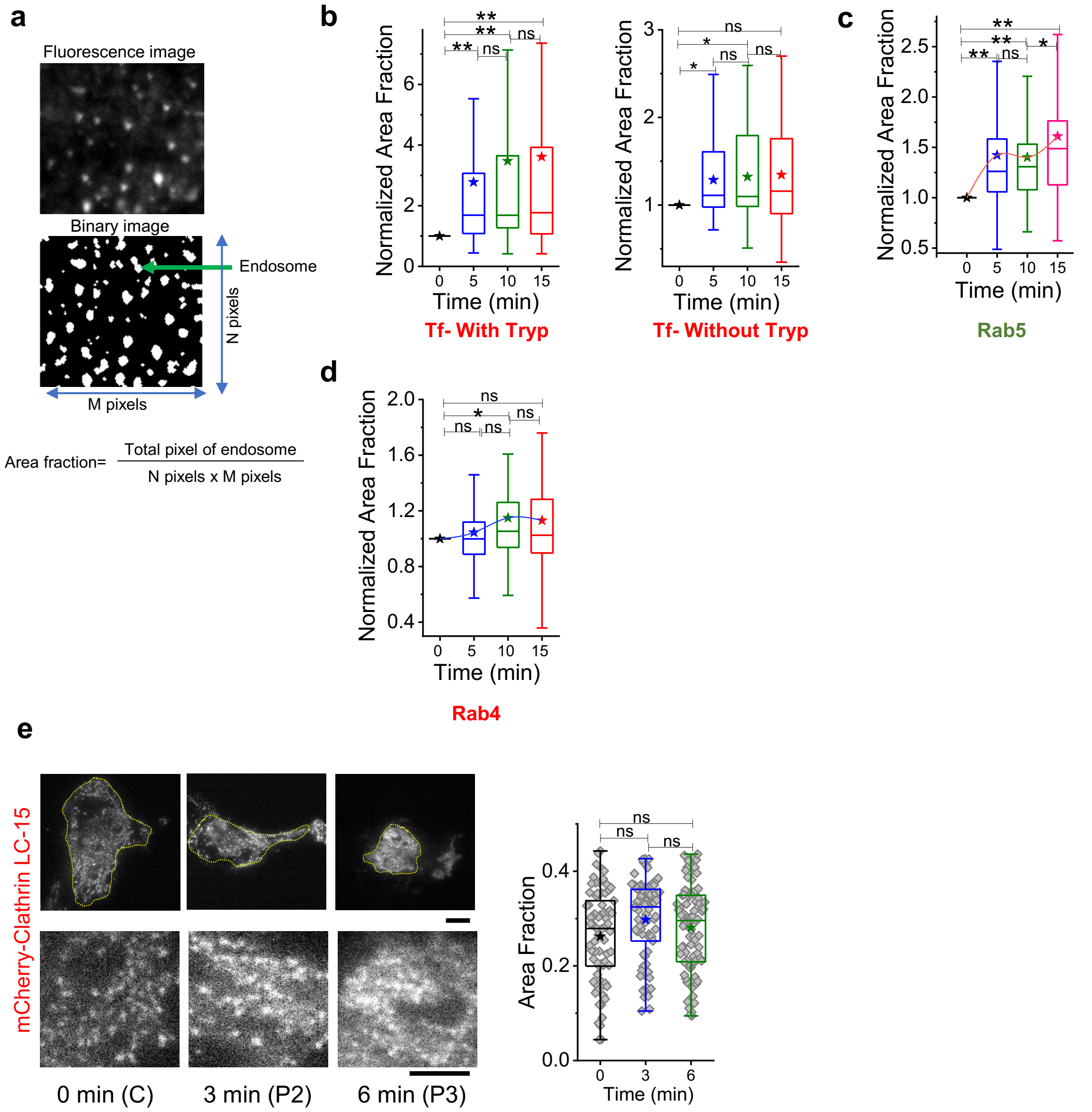

***Figure S6. Fluorescence analysis, Transferrin, Early and Late Endosome imaging, Clathrin puncta analysis. (a)*** Representative fluorescence image and its binary version. Formula depicts how area fraction was calculated. **(b)** *Normalized area fraction (area in $\mu m^2$ covered by Transferrin puncta per $\mu m^2$) of live cells followed over time as spread area reduces on de-adhesion(left) and without adding de-adhering agent (right).* ***(c)*** *Normalized area fraction of Rab 5 of same region followed as the cell de-adheres. Area fraction collated for n=15 regions over n=15 cells. (d) Normalized area fraction of Rab 4 of same region followed as the cell de-adheres.* ***(e)*** *TIRF images of Clathrin LC-15 transfected cells before and after de-adhesion triggered in the sample. Lower columns are zoomed-in images. Area fraction remain non-significant (in minutes) $n_{cell(Control)(0\ min)}$= 74 cells, $n_{cell(control)(3\ min)}$= 82 cells, $n_{cell(control)(6\ min)}$= 94 cells ,Scale bar= 10 µm.*

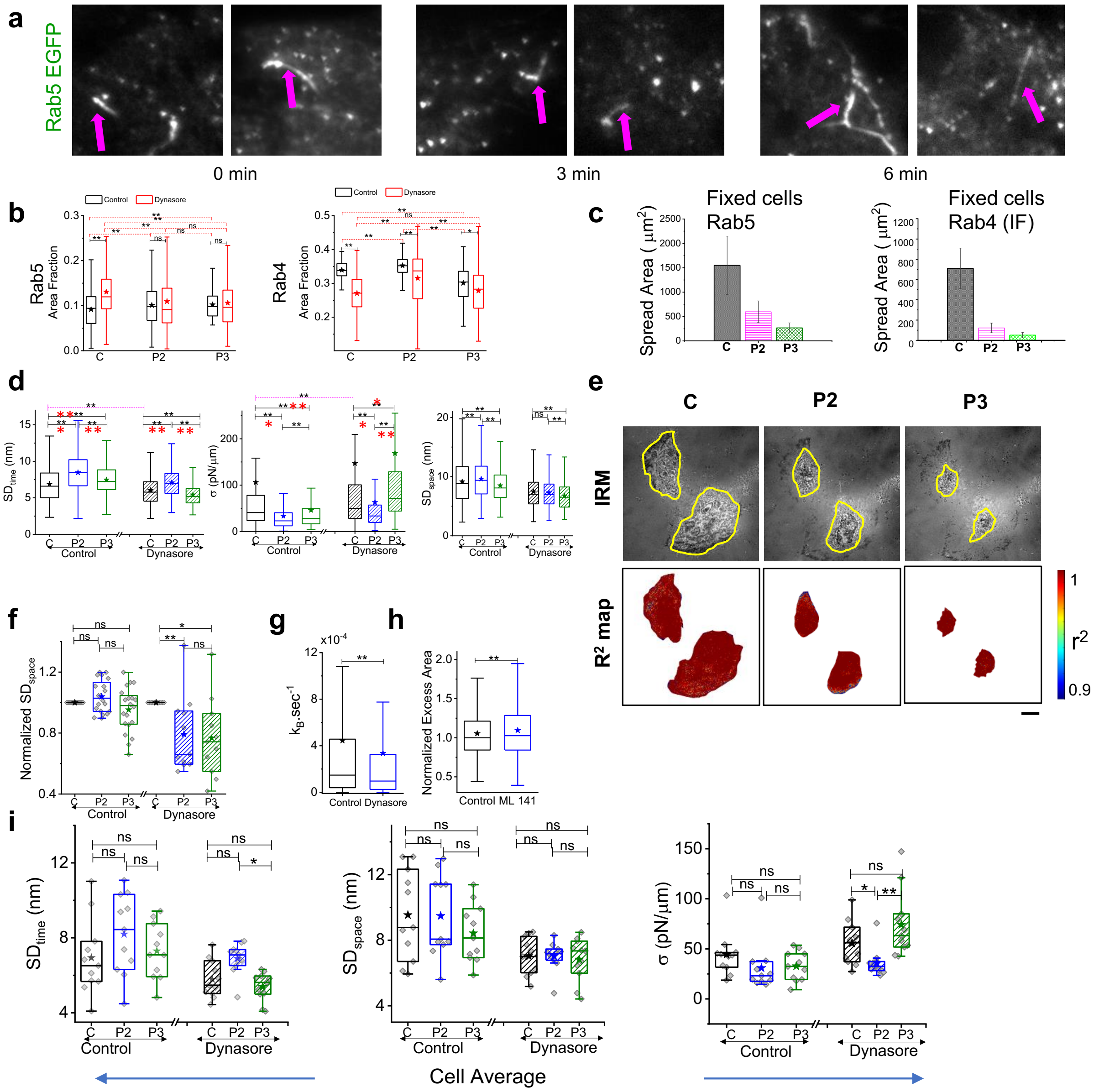

***Figure S7. Early and Late Endosome and Membrane Parameters on blocking endocytosis. (a)*** *Representative images of Rab 5-EGFP in the first two time points after faster de-adhesion in Dynasore-treated cells. Scale bar represents 10 µm.* ***(b)*** *Comparison between area fraction of control and Dynasore treated Rab 5 marked early endosomes (left) and Rab 4 marked late endosomes (right).* ***(c)*** *Spread area of control cells transfected with Rab 5 (left) and immune-stained with Rab 4 before and after addition of 0.25% of trypsin-EDTA.* ***(d)*** *Comparisons made with all FBRs pooled from all cells. n=3 independent experiments. Number of cells=20 (Control), 11 (Dynasore); FBRs of sizes 0.75 µm$^2$ and 4.67 µm$^2$ were used for control and 0.75 µm$^2$ for Dynasore-treated cells.* ***(e)*** *Typical IRM images of Dynasore-treated cells in the different phases of de-adhesion and the corresponding $R^2$ map.* ***(f)*** *Fold change in spatial fluctuations comparing each cell with its own measurements at different phases.* ***(g)*** *Box plots of activity between Control and Dynasore-treated cells using FBR size of 0.75 µm$^2$.* ***(h)*** *Box plots of other normalized excess area between Control and ML 141 treated cells using FBR size of 0.75 µm$^2$.* ***(i)*** *cell-wise comparison of fluctuations and tension. One-way Anova with Bonferroni correction is performed for $SD_{time}$ and $SD_{space}$ since the data is normal. For tension, the Mann Whitney U test is performed and * denotes p value < 0.016 (adjusted by group size of 3 per experiment). n=3 independent experiments. Red * denote LMM was performed.*

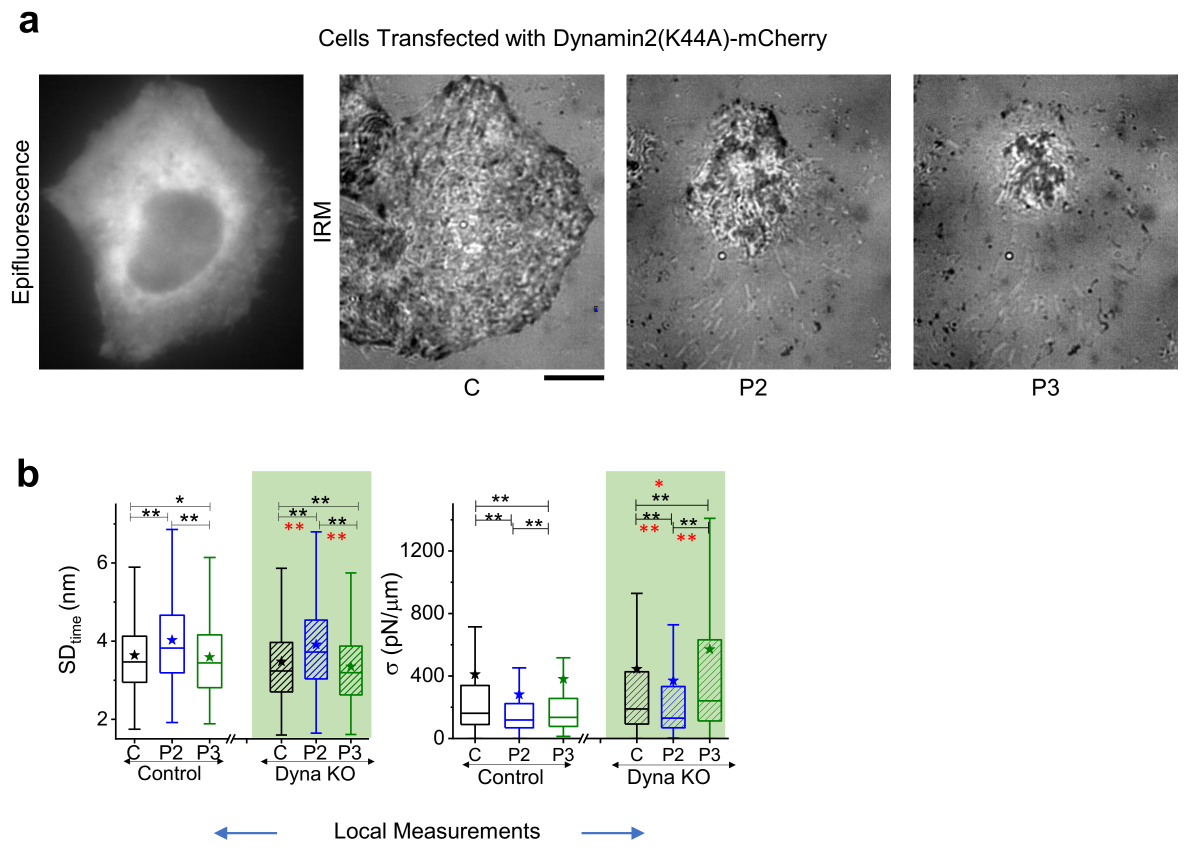

***Figure S8. Fluctuations and Tension of Dynamin2(K44A)-mCherry transfected cell (a)****Representative Epifluorescence and IRM images of HeLa cell transfected with Dynamin2(K44A)-mCherry plasmid. Scale bar = 10 μm.* ***(b)*** *FBR wise comparison of temporal fluctuations and tension.* Red * denote LMM was performed.

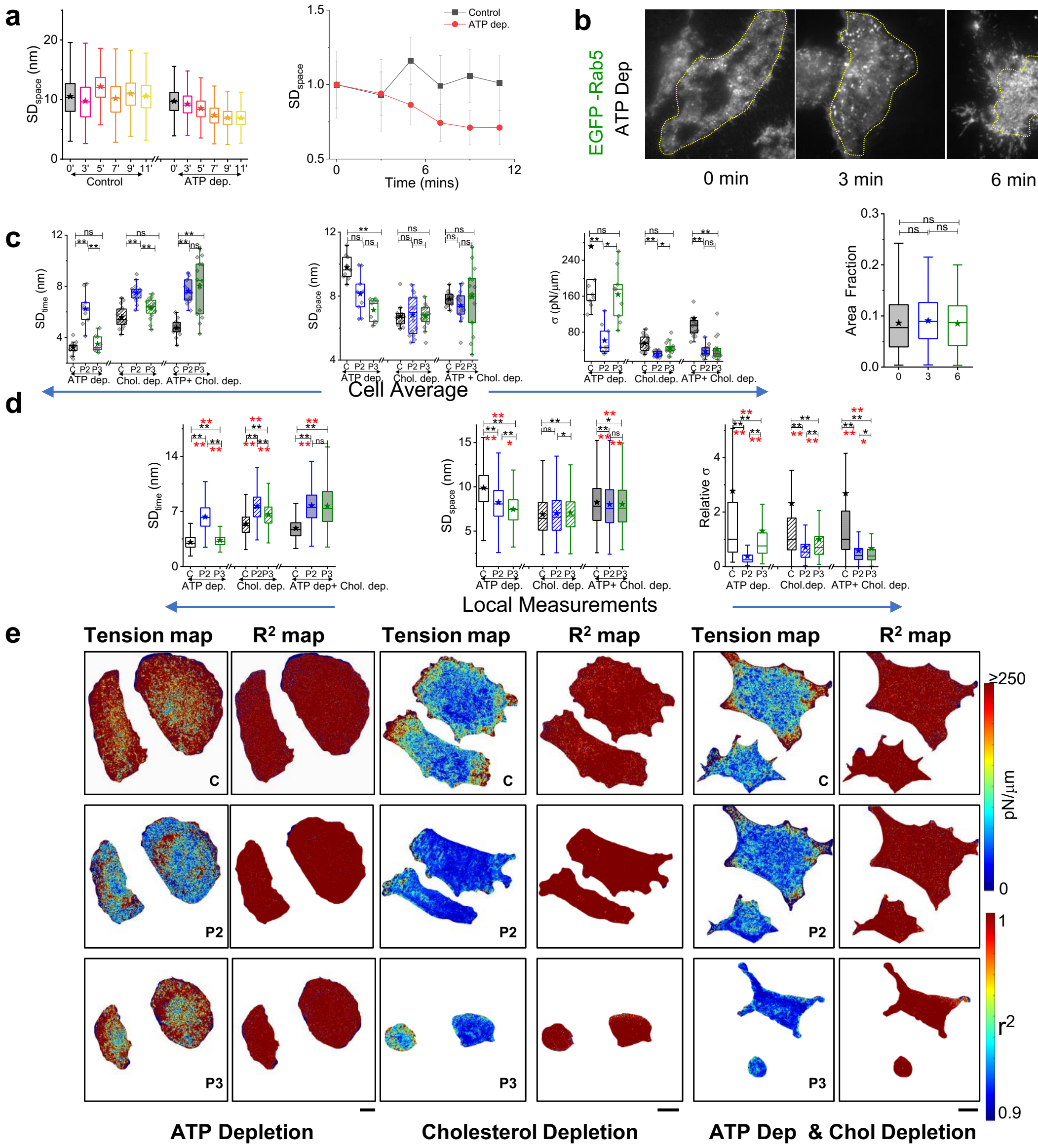

***Figure S9. Membrane fluctuation, Rab 5 area in ATP Depletion, Parameters and tension maps on ATP and cholesterol depletion and both ATP – Cholesterol depletion. (a)*** *Time series boxplots and median (with MAD as error bar, lower panel) for different parameters for control and ATP-depleted cells on de-adhesion using an FBR size of 4.67 $\mu m^2$.* ***(b)*** *TIRF images of different cells transfected with Rab 5 before and after de-adhesion triggered in ATP Depleted cells and Area Fraction of Rab 5****(c)*** *Cell-wise comparison of fluctuations and tension.* ***(d)*** *FBR-wise comparison of fluctuations, relative tension at different phases of de-adhesion in ATP-depleted, cholesterol depleted and ATP-and-cholesterol depleted conditions.* ***(e)*** *Typical tension maps in ATP-depleted, cholesterol depleted and ATP-and-cholesterol depleted conditions at different phases of de-adhesion. Red * denote LMM was performed. Scale bar represents 10 $\mu m$.*

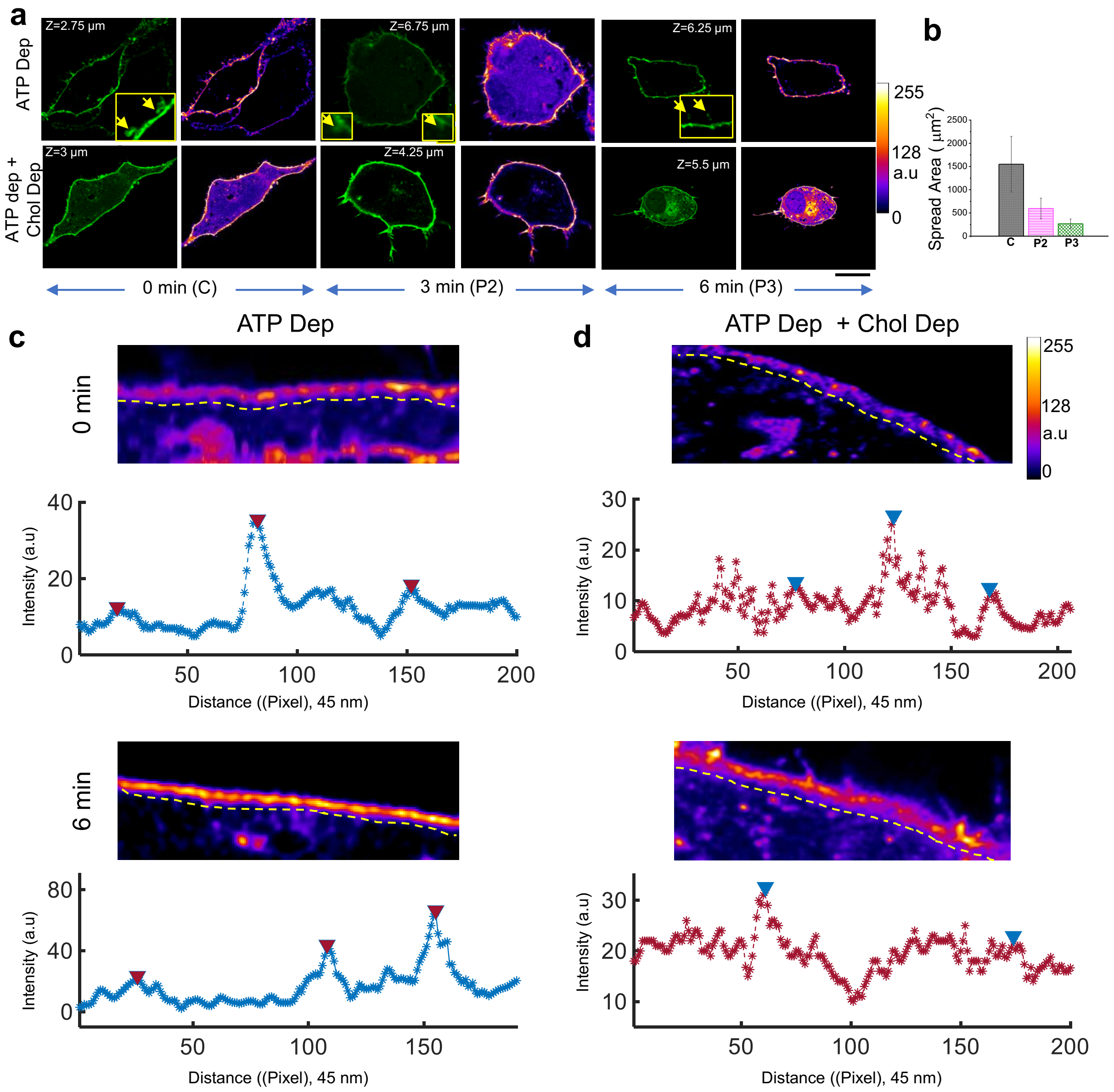

***Figure S10**. **Membrane images marked with EGFP-CAAX.** **(a)** Representative grey scale and colour-coded confocal images of cells (ATP-depleted and ATP Dep+cholesterol-depleted). The rightmost column shows a zoomed-in section highlighting more internal structures in ATP-depleted cells than in ATP Dep +cholesterol-depleted ones. Scale bar = 10 $\mu$m. **(b)** Spread area of CAAX- GFP transfected ATP Depleted cell before and after addition of 0.25% Trypsin- EDTA. Typical images of cell sections showing line ROIs where scans were performed parallel to the membrane in the **(c)** cytosolic side of an ATP depleted cell and **(d)** an ATP Dep +cholesterol-depleted cell at 0 and 6 min after de-adhesion. Plots show intensity line scans with triangles pointing out detected peaks with a minimal width and height. Scale bar = 5 $\mu$m.*

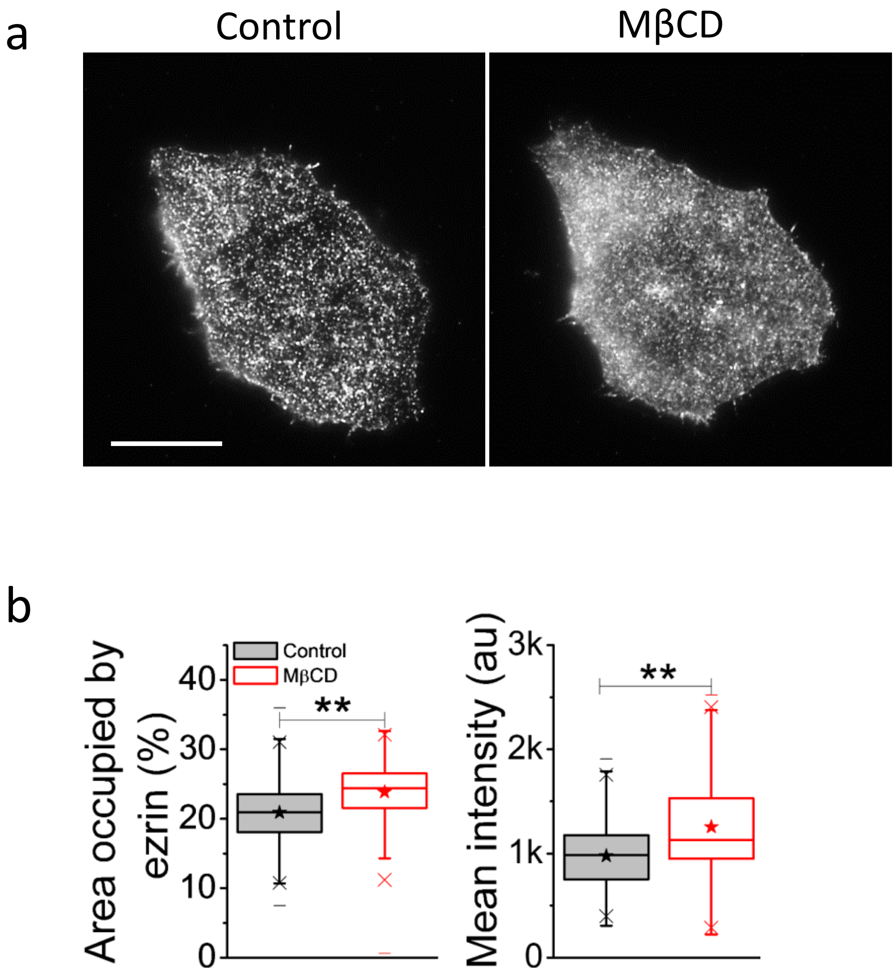

***Figure S11. Enhanced surface ezrin on cholesterol depletion. (a)*** *Representative TIRF images of ezrin immune-stained cell under control and cholesterol-depleted conditions. Scale bar = 10 μm.* ***(b)*** *Boxplots of (left) percentage area and (right) mean intensity occupied by ezrin fluorescent punctas in 3.9 μm x 3.9 μm regions in cells.* $N_{cell}$ *= 15 each.*

| Table S1 |
|---|

List of statistical parameters for all figures

| Fig 1c | | | | | | | | | |
|---|---|---|---|---|---|---|---|---|---|
| | **Conditions** | **$N_{cell}$** | | **Mean** | **SD** | **SEM** | **Median** | **p-values (wrt Faster)** | |
| Time (min) | Control Faster | 19 | | 8.87 | 8.05 | 2.08 | 6.20 | | |
| | Dyna | 7 | | 5.79 | 4.07 | 0.74 | 5.35 | 0.38 | |
| | Cyto D | 10 | | 8.04 | 2.58 | 0.91 | 8.25 | 3.95E-06 | |
| | ATP Dep | 22 | | 1.19 | 1.10 | 0.27 | 0.70 | 0.84 | |
| | Chol Dep | 14 | | 6.06 | 1.47 | 0.33 | 5.75 | 1.30E-05 | |
| | ATP+ Chol Dep | 14 | | 2.20 | 1.16 | 0.27 | 2.05 | 2.23E-01 | |
| | **Conditions** | **$N_{cell}$** | | **Mean** | **SD** | **SEM** | **Median** | **p-values ( wrt slower 0.05% Trypsin)** | |
| Time | Control Slower | 11 | | 12.75 | 7.15 | 2.15 | 10.40 | | |
| | Cyto D Slower | 18 | | 12.18 | 10.40 | 2.45 | 7.25 | 0.38 | |
| **Fig 2b** | | | | | | | | | |
| **Parameters** | **Conditions** | **$N_{cell}$** | **$n_{FBR}$** | **Mean** | **SD** | **SEM** | **Median** | **p-values (wrt C)** | **p-values (wrt P2)** |
| Normalized $SD_{time}$ | C | 22 | | 1.00 | 0.00 | 0.00 | 1.00 | | 1.17E-04 |
| | P2 | 22 | | 1.16 | 0.17 | 0.04 | 1.13 | 1.17E-04 | |
| | P3 | 22 | | 1.09 | 0.19 | 0.04 | 1.10 | 0.03 | 0.28 |
| Normalized Tension | C | 22 | | 1.00 | 0.00 | 0.00 | 1.00 | | 9.69E-04 |
| | P2 | 22 | | 0.75 | 0.28 | 0.06 | 0.68 | 9.69E-04 | |
| | P3 | 22 | | 0.85 | 0.43 | 0.09 | 0.74 | 9.69E-04 | 0.58 |
| **Fig 2f** | | | | | | | | | |
| **Parameters** | **Conditions** | **$N_{cell}$** | **$n_{FBR}$** | **Mean** | **SD** | **SEM** | **Median** | **p-values (wrt C)** | **p-values (wrt P2)** |
| Normaliz | C (Control Slow) | 18 | | 1 | 0 | 0 | 1 | | 2.47E-03 |

| | | | | | | | | | |
|---|---|---|---|---|---|---|---|---|---|
| ed $SD_{time}$ (Cell-wise) | P2 (Control Slow) | 18 | | 1.16 | 0.11 | 0.03 | 1.17 | 2.50E-03 | |
| | P3 (Control Slow) | 18 | | 1.18 | 0.14 | 0.03 | 1.2 | <1E-5 | 0.49614 |
| | C (Cyto D) | 25 | | 1 | 0 | 0 | 1 | | 1.10E-05 |
| | P2 (Cyto D) | 25 | | 1.1 | 0.12 | 0.02 | 1.09 | 1.10E-05 | |
| | P3 (Cyto D) | 25 | | 1.04 | 0.1 | 0.02 | 1.03 | 7.10E-02 | 0.06817 |
| Normalized Tension (Cell-wise) | C (Control Slow) | 18 | | 1 | 0 | 0 | 1 | | 2.78E-04 |
| | P2 (Control Slow) | 18 | | 0.75 | 0.21 | 0.05 | 0.71 | 2.80E-04 | |
| | P3 (Control Slow) | 18 | | 0.67 | 0.16 | 0.04 | 0.69 | <1E-5 | 0.33456 |
| | C (Cyto D) | 25 | | 1 | 0 | 0 | 1 | | 0.44 |
| | P2 (Cyto D) | 25 | | 0.96 | 0.27 | 0.05 | 0.92 | 0.44 | |
| | P3 (Cyto D) | 25 | | 0.99 | 0.27 | 0.05 | 0.98 | 0.07 | 0.94 |
| **Fig 2g** | | | | | | | | | |
| **Parameters** | **Conditions** | **$N_{cell}$** | **$n_{FBR}$** | **Mean** | **SD** | **SEM** | **Median** | **p-values (wrt Control)** | |
| Force (pN) | Control | 10 | | 12.87 | 4.87 | 1.54 | 11.23 | | |
| | Trypsin | 10 | | 10.73 | 2.91 | 0.92 | 9.9 | 0.4055 | |
| **Fig 4e** | | | | | | | | | |
| **Parameters** | **Conditions** | **$N_{cell}$** | **$n_{FBR}$** | **Mean** | **SD** | **SEM** | **Median** | **p-values (wrt C)** | **p-values (wrt P2)** |
| Tension (pN/µm) | C (Control) | 20 | 2006 | 106.1 | 266.6 | 5.95 | 40.7 | | 0 |
| | P2 (Control) | 20 | 957 | 33.05 | 35.41 | 1.14 | 22.79 | 0 | |
| | P3 (Control) | 20 | 355 | 45.8 | 82.57 | 4.38 | 27.76 | <1E-5 | <1E-5 |
| | C (Dynasore) | 11 | 6669 | 147.1 | 358.3 | 4.37 | 49.67 | | 0 |
| | P2 (Dynasore) | 11 | 3230 | 62.89 | 156.3 | 2.75 | 33.42 | 0 | |

| | | | | | | | | | |
|---|---|---|---|---|---|---|---|---|---|
| | P3 (Dynasore) | 11 | 1475 | 168.6 | 326.9 | 8.51 | 70.99 | <1E-5 | <1E-5 |
| **Fig 4f** | | | | | | | | | |
| $SD_{time}$ (nm) | C (Control) | 20 | 3184 | 6.88 | 2.43 | 0.04 | 6.59 | | <1E-5 |
| | P2 (Control) | 20 | 1859 | 8.47 | 2.56 | 0.06 | 8.44 | <1E-5 | |
| | P3 (Control) | 20 | 796 | 7.47 | 2.03 | 0.07 | 7.23 | <1E-5 | 0 |
| | C (Dynasore) | 11 | 8861 | 6.01 | 1.99 | 0.02 | | | <1E-5 |
| | P2 (Dynasore) | 11 | 4042 | 7.08 | 2.02 | 0.03 | 6.98 | <1E-5 | |
| | P3 (Dynasore) | 11 | 1733 | 5.42 | 1.57 | 0.04 | 5.17 | 0 | 0 |
| **Fig 4h** | | | | | | | | | |
| **Parameters** | **Conditions** | **$N_{cell}$** | **$n_{FBR}$** | **Mean** | **SD** | **SEM** | **Median** | **p-values (wrt Control)** | |
| Normalized Excess Area | Control | 15 | 3741 | 1.07 | 0.42 | 0.01 | 1 | | |
| | Dynasore | 11 | 6669 | 0.99 | 0.35 | 0.00 | 0.9 | 0 | |
| **Fig 5c** | | | | | | | | | |
| **Parameters** | **Conditions** | **$N_{cell}$** | **$n_{patches}$** | **Mean** | **SD** | **SEM** | **Median** | **p-value (wrt 0 min )** | **p-value (wrt 3 min)** |
| Area Fraction Rab5(ATP Dep) | 0 min | 26 | 168 | 0.09 | 0.06 | 0.005 | 0.08 | | 0.07 |
| | 3 min | 41 | 288 | 0.09 | 0.05 | 0.003 | 0.09 | 0.07 | |
| | 6min | 39 | 255 | 0.08 | 0.05 | 0.003 | 0.09 | 0.52 | 0.17 |
| **Fig 6c** | | | | | | | | | |
| **Parameters** | **Conditions** | **$N_{cell}$** | **$n_{FBR}$** | **Mean** | **SD** | **SEM** | **Median** | **p-values (wrt C)** | **p-values (wrt P2)** |
| Normalized $SD_{time}$ | C (ATP Dep.) | 9 | | 1 | 0 | 0 | 1 | | 1.61E-04 |
| | P2 (ATP Dep.) | 9 | | 1.99 | 0.46 | 0.15 | 2.08 | 1.60E-04 | |
| | P3 (ATP Dep.) | 9 | | 1.11 | 0.23 | 0.08 | 1.19 | 0.22 | 0.002 |

| | | | | | | | | | |
|---|---|---|---|---|---|---|---|---|---|
| (Cell-wise) | C (Chol. Dep.) | 16 | | 1 | 0 | 0 | 1 | | <1E-5 |
| | P2 (Chol. Dep.) | 16 | | 1.36 | 0.19 | 0.05 | 1.33 | <1E-5 | |
| | P3 (Chol. Dep.) | 16 | | 1.14 | 0.19 | 0.05 | 1.11 | 0.001 | 0.002 |
| | C (ATP Dep. +Chol. Dep.) | 15 | | 1 | 0 | 0 | 1 | | <1E-5 |
| | P2 (ATP Dep. +Chol. Dep.) | 15 | | 1.63 | 0.27 | 0.07 | 1.53 | <1E-5 | |
| | P3 (ATP Dep. +Chol. Dep.) | 15 | | 1.69 | 0.43 | 0.11 | 1.75 | 1.70E-05 | 0.45 |
| Normalized Tension (Cell-wise) | C (ATP Dep.) | 9 | | 1 | 0 | 0 | 1 | | 1.61E-04 |
| | P2 (ATP Dep.) | 9 | | 0.29 | 0.21 | 0.07 | 0.23 | 1.60E-04 | |
| | P3 (ATP Dep.) | 9 | | 0.79 | 0.35 | 0.12 | 0.83 | 0.22 | 0.002 |
| | C (Chol. Dep.) | 16 | | 1 | 0 | 0 | 1 | | <1E-5 |
| | P2 (Chol. Dep.) | 16 | | 0.62 | 0.15 | 0.04 | 0.59 | <1E-5 | |
| | P3 (Chol. Dep.) | 16 | | 0.85 | 0.25 | 0.06 | 0.87 | 0.05 | 0.006 |
| | C (ATP Dep. +Chol. Dep.) | 15 | | 1 | 0 | 0 | 1 | | <1E-5 |
| | P2 (ATP Dep. +Chol. Dep.) | 15 | | 0.39 | 0.15 | 0.04 | 0.37 | <1E-5 | |
| | P3 (ATP Dep. +Chol. Dep.) | 15 | | 0.44 | 0.37 | 0.09 | 0.34 | 1.71E-05 | 0.65 |

| Fig 6e | | | | | | | | | |
|---|---|---|---|---|---|---|---|---|---|
| **Parameters** | **Conditions** | **$N_{cell}$** | **$n_{FBR}$** | **Mean** | **SD** | **SEM** | **Median** | **p-values (wrt Normal Faster)** | **Signification from 0.25%** |

| | | | | | | | | | |
|---|---|---|---|---|---|---|---|---|---|
| | Normal Faster (0.25%) | 22 | | -0.03 | 0.08 | 0.02 | -0.05 | | 0.06 |
| C to P2 (Cell-wise) | Dynasore | 11 | | -0.05 | 0.06 | 0.02 | -0.07 | 0.83 | 1.41E-02 |
| | ATP Dep | 9 | | -0.12 | 0.06 | 0.02 | -0.11 | 0.00 | 4.14E-04 |
| | Chol Dep | 16 | | -0.21 | 0.11 | 0.03 | -0.18 | 9.35E-07 | <1E-5 |
| | ATP+Chol Dep | 15 | | -0.18 | 0.07 | 0.02 | -0.18 | 4.37E-06 | <1E-5 |
| P3 to P2 (Cell-wise) | Normal Faster(0.25%) | 22 | | 0.05 | 0.16 | 0.03 | 0.03 | | 0.15 |
| | Dynasore | 11 | | 0.17 | 0.15 | 0.05 | 0.14 | 0.02 | 0 |
| | ATP Dep | 9 | | 0.15 | 0.09 | 0.03 | 0.15 | 0.01 | 9.12E-04 |
| | Chol Dep | 16 | | 0.12 | 0.1 | 0.03 | 0.12 | 0.01 | 2.53E-04 |
| | ATP+Chol Dep | 15 | | 0.02 | 0.21 | 0.05 | -0.03 | 0.09 | 0.71 |
| P3 to C (Cell-wise) | Normal Faster(0.25%) | 22 | | -0.01 | 0.06 | 0.01 | | -0.02 | 0.50 |
| | Dynasore | 11 | | 0.04 | 0.05 | 0.02 | 0.03 | 0.01 | 0.04 |
| | ATP Dep | 9 | | -0.01 | 0.02 | 0.01 | -0.01 | 0.53 | 0.11 |
| | Chol Dep | 16 | | -0.02 | 0.04 | 0.01 | -0.03 | 0.94 | 0.07 |
| | ATP+Chol Dep | 15 | | -0.07 | 0.06 | 0.02 | -0.07 | 2.23E-05 | 5.82E-04 |

| Fig 7d | | | | | | | | | |
|---|---|---|---|---|---|---|---|---|---|
| **Parameters** | **Conditions** | **$N_{cell}$** | **$n_{ROI}$** | **Mean** | **SD** | **SEM** | **Median** | **p-values (wrt 0 min)** | **p-values (wrt 3 min)** |
| No of tubules ($\mu m^{-1}$) | ATP Dep (0 min) | 20 | 139 | 0.45 | 0.25 | 0.02 | 0.43 | | <1E-5 |
| | ATP Dep (3 min) | 18 | 124 | 0.58 | 0.25 | 0.02 | 0.59 | <1E-5 | |
| | ATP Dep (6 min) | 28 | 240 | 0.49 | 0.23 | 0.01 | 0.47 | 0.14 | 3.61E-05 |
| | ATP +Chol Depletion (0 min) | 21 | 149 | 0.49 | 0.21 | 0.02 | 0.48 | | 9.46E-03 |
| | ATP +Chol Depletion (3 min) | 24 | 161 | 0.41 | 0.27 | 0.02 | 0.39 | 9.46E-03 | |

| | | | | | | | | | |
|---|---|---|---|---|---|---|---|---|---|
| | ATP +Chol Depletion (6 min) | 21 | 80 | 0.41 | 0.21 | 0.02 | 0.38 | 0.004 | 6.91E-01 |
| **Fig 7e** | | | | | | | | | |
| **Parameters** | **Conditions** | **$N_{cell}$** | **$n_{ROI}$** | **Mean** | **SD** | **SEM** | **Median** | **p-values (wrt 0 min)** | **p-values (wrt 3 min)** |
| Intensity (a.u) | ATP Dep (0 min) | 20 | 16108 | 16.91 | 10.56 | 0.08 | 14.56 | | 0 |
| | ATP Dep (3 min) | 18 | 18586 | 35.89 | 17.51 | 0.13 | 32.06 | 0 | |
| | ATP Dep (6 min) | 28 | 28062 | 26.06 | 15.52 | 0.09 | 22.96 | 0 | 0 |
| | ATP +Chol Depletion (0 min) | 21 | 15301 | 11.68 | 8.67 | 0.07 | 10 | | 0 |
| | ATP +Chol Depletion (3 min) | 24 | 5166 | 19.72 | 6.68 | 0.09 | 19.68 | 0 | |
| | ATP +Chol Depletion (6 min) | 21 | 11615 | 17.40 | 9.31 | 0.09 | 16.34 | 0 | 0 |

**Supplementary Tables**

| | | | | | | | | | | |
|---|---|---|---|---|---|---|---|---|---|---|
| **Fig S1b** | | | | | | | | | | |
| **Parameters** | **Conditions** | **$N_{cell}$** | **$n_{FBR}$** | **Mean** | **SD** | **SEM** | **Median** | **p-values (wrt 1min )** | | |
| $SD_{time}$(nm) | 1 min | 5 | 1102 | 5.8 | 1.2 | 0.04 | 5.8 | | | |
| | 20 min | 5 | 1207 | 5.7 | 1.2 | 0.03 | 5.7 | 0.14 | | |
| $SD_{space}$ (nm) | 1 min | 5 | 1102 | 8.1 | 2.3 | 0.07 | 7.9 | | | |
| | 20 min | 5 | 1207 | 8.2 | 2.3 | 0.07 | 8 | 0.26 | | |
| **Fig S1c** | | | | | | | | | | |
| σ | 1 min | 5 | 444 | 299.15 | 534.45 | 25.36 | 73.67 | | | |
| | 20 min | 5 | 519 | 328.58 | 600.38 | 26.35 | 83.01 | 0.48 | | |
| **Fig S2b** | | | | | | | | | | |
| **Parameters** | **Conditions** | **$N_{cell}$** | **$n_{FBR}$** | **Mean** | **SD** | **SEM** | **Median** | **p-values (wrt C)** | **p-values (wrt P1)** | **p-values (wrt P2)** |
| $SD_{time}$ (nm) | C (Slower) | 6 | 287 | 3.34 | 0.66 | 0.04 | 3.27 | | 0.14 | $<10^{-5}$ |
| | P1 (Slower) | 6 | 287 | 3.4 | 0.62 | 0.04 | 3.39 | 0.14 | | $<10^{-5}$ |

| | | | | | | | | | | |
|---|---|---|---|---|---|---|---|---|---|---|
| | P2 (Slower) | 6 | 287 | 3.76 | 0.74 | 0.04 | 3.69 | $<10^{-5}$ | $<10^{-5}$ | |
| | P3 (Slower) | 6 | 287 | 4.05 | 0.76 | 0.04 | 3.94 | $<10^{-5}$ | $<10^{-5}$ | $<10^{-5}$ |
| | C (Faster) | 5 | 155 | 3.81 | 0.95 | 0.08 | 3.79 | | | $<10^{-5}$ |
| | P1 (Faster) | 5 | 155 | 4.05 | 0.81 | 0.07 | 4.03 | 0.013 | 0.013 | $<10^{-5}$ |
| | P2 (Faster) | 5 | 155 | 4.47 | 0.76 | 0.06 | 4.54 | $<10^{-5}$ | $<10^{-5}$ | |
| | P3 (Faster) | 5 | 155 | 4.19 | 0.87 | 0.07 | 4.07 | 4.00E-04 | 0.22 | 6.00E-04 |

| **Fig S2c** | | | | | | | | | | |
|---|---|---|---|---|---|---|---|---|---|---|
| $SD_{space}$ (nm) | C (Slower) | 6 | 287 | 5.11 | 1.35 | 0.07 | 4.91 | | 0.49 | 0.45 |
| | P1 (Slower) | 6 | 287 | 5.19 | 1.41 | 0.08 | 5.12 | 0.49 | | 0.93 |
| | P2 (Slower) | 6 | 287 | 5.17 | 1.25 | 0.07 | 5.07 | 0.45 | 0.93 | |
| | P3 (Slower) | 6 | 287 | 4.87 | 1.22 | 0.06 | 4.77 | 3.00E-02 | 4.00E-03 | 2.00E-03 |
| | C (Faster) | 5 | 155 | 5.41 | 1.3 | 0.1 | 5.34 | | 0.004 | 0.76 |
| | P1 (Faster) | 5 | 155 | 5.89 | 1.37 | 0.11 | 5.79 | 0.004 | | 9.00E-03 |
| | P2 (Faster) | 5 | 155 | 5.5 | 1.34 | 0.11 | 5.36 | 0.76 | 9.00E-03 | |
| | P3 (Faster) | 5 | 155 | 5.06 | 1.52 | 0.12 | 4.78 | 0.003 | $<10^{-5}$ | 7.00E-04 |
| **Fig S2d** | | | | | | | | | | |
| **Parameter** | **Conditions** | **$N_{cell}$** | **$n_{FBR}$** | **Mean** | **SD** | **SEM** | **Median** | **p-values (wrt C)** | **p-values (wrt P1)** | **p-values (wrt P2)** |
| Tension (pN/µm) | C (Slower) | 6 | 184 | 335.18 | 509.56 | 37.57 | 149.47 | | 0.86 | 1.00E-04 |
| | P1 (Slower) | 6 | 202 | 362.72 | 594.72 | 41.84 | 136.41 | 0.86 | | 3.00E-04 |
| | P2 (Slower) | 6 | 186 | 246.62 | 418.89 | 30.71 | 95.09 | 1.00E-04 | 3.00E-04 | |
| | P3 (Slower) | 6 | 189 | 222.29 | 388.63 | 28.27 | 78.86 | $<10^{-5}$ | $<10^{-5}$ | 0.16 |
| | C (Faster) | 5 | 75 | 360.65 | 672.46 | 77.65 | 96.22 | | 0.17 | 0.01 |
| | P1 (Faster) | 5 | 72 | 164.65 | 256.43 | 30.22 | 75.81 | 0.17 | | 0.13 |
| | P2 (Faster) | 5 | 73 | 231 | 380.72 | 44.56 | 60.85 | 0.01 | 0.13 | |
| | P3 (Faster) | 5 | 88 | 314.63 | 499.88 | 53.29 | 118.08 | 0.7 | 0.08 | 0.01 |

| Fig S3b | | | | | | | | | |
|---|---|---|---|---|---|---|---|---|---|
| **Parameters** | **Conditions** | **$N_{cell}$** | **$n_{FBR}$** | **Mean** | **SD** | **SEM** | **Median** | **p-values (wrt C)** | **p-values (wrt P2)** |
| $SD_{time}$ (nm) | C (TrypLE) | 24 | 19377 | 5.96 | 1.72 | 0.01 | 5.73 | | 0 |
| | P2 (TrypLE) | 24 | 10561 | 7.02 | 1.87 | 0.02 | 6.8 | 0 | |
| | P3 (TrypLE)) | 24 | 2819 | 6.52 | 1.36 | 0.03 | 6.37 | $<10^{-5}$ | 0 |
| $SD_{space}$ (nm) | C (TrypLE) | 24 | 19377 | 6.91 | 2.18 | 0.02 | 6.6 | | $<10^{-5}$ |
| | P2 (TrypLE) | 24 | 10561 | 7.39 | 2.51 | 0.02 | 6.93 | $<10^{-5}$ | |
| | P3 (TrypLE) | 24 | 2819 | 6.44 | 1.82 | 0.03 | 6.21 | 0 | 0 |
| Ten. (pN/µm) | C (TrypLE) | 24 | 14038 | 98.88 | 242.49 | 2.05 | 47.25 | | 0 |
| | P2 (TrypLE) | 24 | 6717 | 49.87 | 109.27 | 1.33 | 32.28 | 0 | |
| | P3 (TrypLE)) | 24 | 1178 | 56.57 | 84.22 | 2.45 | 40.38 | $<10^{-5}$ | $<10^{-5}$ |
| **Fig S3c** | | | | | | | | | |
| **Parameters** | **Conditions** | **$N_{cell}$** | **$n_{FBR}$** | **Mean** | **SD** | **SEM** | **Median** | **p-values (wrt C)** | **p-values (wrt P2)** |
| | C (Slower) | 18 | | 1 | 0 | 0 | 1 | | 0.002 |
| Normalized $SD_{time}$ | P2 (Slower) | 18 | | 1.16 | 0.11 | 0.03 | 1.17 | 0.002 | |
| | P3 (Slower) | 18 | | 1.18 | 0.14 | 0.03 | 1.2 | <1E-5 | 0.5 |
| | C (Faster) | 22 | | 1.00 | 0.00 | 0.00 | 1 | | 1.17E-04 |
| | P2 (Faster) | 22 | | 1.16 | 0.17 | 0.04 | 1.13 | 1.17E-04 | |
| | P3 (Faster) | 22 | | 1.09 | 0.19 | 0.04 | 1.10 | 0.03 | 0.28 |
| Normalized $Sd_{space}$ | C (Slower) | 18 | | 1 | 0 | 0 | 1 | | 2.18E-05 |
| | P2 (Slower) | 18 | | 1.07 | 0.05 | 0.01 | 1.07 | 2.18E-05 | |
| | P3 (Slower) | 18 | | 1.05 | 0.1 | 0.02 | 1.06 | 0.02 | 0.49614 |
| | C (Faster) | 22 | | 1 | 0 | 0 | 1 | | 1 |
| | P2 (Faster) | 22 | | 1.02 | 0.11 | 0.02 | 1.00 | 1 | |
| | P3 (Faster) | 22 | | 0.94 | 0.15 | 0.03 | 0.95 | 0.10 | 0.06 |

| | | | | | | | | | |
|---|---|---|---|---|---|---|---|---|---|
| Normalized Tension | C (Slower) | 18 | | 1.00 | 0.00 | 0.00 | 1.00 | | 2.78E-04 |
| | P2 (Slower) | 18 | | 0.75 | 0.21 | 0.05 | 0.71 | 2.78E-04 | |
| | P3 (Slower) | 18 | | 0.67 | 0.16 | 0.04 | 0.69 | <1E-5 | 0.33 |
| | C (Faster) | 22 | | 1.00 | 0.00 | 0.00 | 1.00 | | 9.69E-04 |
| | P2 (Faster) | 22 | | 0.75 | 0.28 | 0.06 | 0.68 | 9.69E-04 | |
| | P3 (Faster) | 22 | | 0.85 | 0.43 | 0.09 | 0.74 | 9.69E-04 | 0.58122 |

| Fig S4a | | | | | | | | | |
|---|---|---|---|---|---|---|---|---|---|
| **Parameters** | **Conditions** | **$N_{cell}$** | **$n_{FBR}$** | **Mean** | **SD** | **SEM** | **Median** | **p-values (wrt C)** | **p-values (wrt P2)** |
| $SD_{time}$ (nm) | C (Control) | 18 | | 5.12 | 0.55 | 0.13 | 5.23 | | 0.002 |
| | P2 (Control) | 18 | | 5.89 | 0.68 | 0.16 | 5.59 | 0.002 | |
| | P3 (Control) | 18 | | 6.02 | 0.78 | 0.18 | 5.93 | 0.001 | 0.65 |
| | C (Cyto D) | 25 | | 5.16 | 0.48 | 0.10 | 5.11 | | 0.004 |
| | P2 (Cyto D) | 25 | | 5.66 | 0.68 | 0.14 | 5.58 | 0.004 | |
| | P3 (Cyto D) | 25 | | 5.36 | 0.63 | 0.13 | 5.35 | 0.19 | 0.06 |
| $SD_{space}$ (nm) | C (Control) | 18 | | 6.03 | 0.51 | 0.12 | 6.05 | | 0.009 |
| | P2 (Control) | 18 | | 6.45 | 0.58 | 0.14 | 6.26 | 0.009 | |
| | P3 (Control) | 18 | | 6.33 | 0.65 | 0.15 | 6.48 | 0.03 | 0.52 |
| | C (Cyto D) | 25 | | 6.51 | 0.62 | 0.12 | 6.48 | | 0.53 |
| | P2 (Cyto D) | 25 | | 6.48 | 0.67 | 0.13 | 6.37 | 0.53 | |
| | P3 (Cyto D) | 25 | | 6.20 | 0.67 | 0.13 | 6.28 | 0.23 | 0.63 |
| Tension (pN/µm) | C (Control) | 18 | | 68.08 | 21.15 | 4.98 | 64.18 | | 0.004 |
| | P2 (Control) | 18 | | 49.78 | 17.30 | 4.08 | 48.84 | 0.004 | |
| | P3 (Control) | 18 | | 44.52 | 15.12 | 3.56 | 38.85 | 3.72E-04 | 0.26 |
| | C (Cyto D) | 25 | | 59.83 | 14.30 | 2.86 | 56.96 | | 0.23 |
| | P2 (Cyto D) | 25 | | 57.23 | 21.19 | 4.24 | 53.76 | 0.23 | |
| | P3 (Cyto D) | 25 | | 58.65 | 21.12 | 4.22 | 55.99 | 0.35 | 0.80 |

| | | | | | | | | | |
|---|---|---|---|---|---|---|---|---|---|
| Normalized $SD_{space}$ (nm) | C (Control) | 18 | | 1 | 0 | 0 | 1 | | 2.18E-05 |
| | P2 (Control) | 18 | | 1.07 | 0.05 | 0.01 | 1.07 | 2.10E-05 | |
| | P3 (Control) | 18 | | 1.05 | 0.1 | 0.02 | 1.06 | 1.60E-02 | 0.49614 |
| | C (Cyto D) | 25 | | 1 | 0 | 0 | 1 | | 0.19847 |
| | P2 (Cyto D) | 25 | | 1 | 0.08 | 0.02 | 0.99 | 0.1985 | |

| | | | | | | | | | |
|---|---|---|---|---|---|---|---|---|---|
| | P3 (Cyto D) | 25 | | 0.96 | 0.11 | 0.02 | 0.97 | 4.50E-03 | 0.15106 |

| Fig S4b | | | | | | | | | |
|---|---|---|---|---|---|---|---|---|---|
| **Parameters** | **Conditions** | **$N_{cell}$** | **$n_{FBR}$** | **Mean** | **SD** | **SEM** | **Median** | **p-values (wrt C)** | **p-values (wrt P2)** |
| $SD_{time}$ (nm) | C (Control Slow) | 18 | 16440 | 5.64 | 1.71 | 0.01 | 5.35 | | <1E-5 |
| | P2 (Control Slow) | 18 | 11475 | 6.12 | 1.91 | 0.02 | 5.85 | <1E-5 | |
| | P3 (Control Slow) | 18 | 4898 | 6.16 | 1.76 | 0.03 | 5.92 | <1E-5 | 0.008 |
| | C (Cyto D) | 25 | 16126 | 5.29 | 1.38 | 0.01 | 5.13 | | <1E-5 |
| | P2 (Cyto D) | 25 | 9693 | 5.82 | 1.29 | 0.01 | 5.69 | <1E-5 | |
| | P3 (Cyto D) | 25 | 6375 | 5.47 | 1.2 | 0.02 | 5.32 | <1E-5 | 0 |
| Relative Tension | C (Control Slow) | 18 | 11900 | 2.52 | 5.45 | 0.05 | 1 | | 0 |
| | P2 (Control Slow) | 18 | 8308 | 1.63 | 3.94 | 0.04 | 0.71 | 0 | |
| | P3 (Control Slow) | 18 | 3096 | 1.24 | 2.65 | 0.05 | 0.66 | 0 | 2.02E-05 |
| | C (Cyto D) | 25 | 10991 | 2.75 | 6.57 | 0.06 | 1 | | <1E-5 |
| | P2 (Cyto D) | 25 | 6727 | 1.57 | 3.09 | 0.04 | 0.92 | <1E-5 | |
| | P3 (Cyto D) | 25 | 4060 | 1.87 | 4.03 | 0.06 | 0.98 | 0.0166 | <1E-5 |
| $SD_{space}$ (nm) | C (Control Slow) | 18 | 16440 | 6.16 | 1.94 | 0.02 | 5.83 | | <1E-5 |
| | P2 (Control Slow) | 18 | 11475 | 6.53 | 2.17 | 0.02 | 6.17 | <1E-5 | |
| | P3 (Control Slow) | 18 | 4898 | 6.49 | 1.97 | 0.03 | 6.23 | <1E-5 | 0.3 |
| | C (Cyto D) | 25 | 16126 | 6.71 | 2.01 | 0.02 | 6.39 | | 2.21E-05 |
| | P2 (Cyto D) | 25 | 9693 | 6.78 | 1.91 | 0.02 | 6.47 | 2.00E-05 | |
| | P3 (Cyto D) | 25 | 6375 | 6.61 | 1.89 | 0.02 | 6.35 | 0.06 | <1E-5 |

| Fig S6b |
|---|

| Parameters | Conditions | $N_{cell}$ | $n_{ROI}$ | Mean | SD | SEM | Median | p-values (wrt 0 min) | p-values (wrt 5 min) |
|---|---|---|---|---|---|---|---|---|---|
| Normalized Area Fraction (Transferrin with trypsin) | 0 min | 53 | 92 | 1.00 | 0.00 | 0.00 | 1.00 | | 2.92E-15 |
| | 5 min | 53 | 92 | 2.78 | 4.92 | 0.51 | 1.69 | 2.92E-15 | |
| | 10 min | 53 | 92 | 3.47 | 6.45 | 0.67 | 1.69 | 2.63E-19 | 0.36 |
| | 15 min | 53 | 92 | 3.61 | 6.00 | 0.63 | 1.77 | 2.92E-15 | |
| **Parameters** | **Conditions** | **$N_{cell}$** | **$n_{ROI}$** | **Mean** | **SD** | **SEM** | **Median** | **p-values (wrt 0 min)** | **p-values (wrt 5 min)** |
| Normalized Area Fraction (Transferrin without trypsin) | 0 min | 16 | 31 | 1.00 | 0.00 | 0.00 | 1.00 | | 2.49E-03 |
| | 5 min | 16 | 31 | 1.29 | 0.45 | 0.08 | 1.11 | 2.49E-03 | |
| | 10 min | 16 | 31 | 1.32 | 0.55 | 0.10 | 1.10 | 2.49E-03 | 0.94 |
| | 15 min | 16 | 31 | 1.34 | 0.66 | 0.12 | 1.16 | 0.49 | |

| Fig S6c | | | | | | | | | |
|---|---|---|---|---|---|---|---|---|---|
| **Parametes** | **Conditions** | **$N_{cell}$** | **$n_{ROI}$** | **Mean** | **SD** | **SEM** | **Median** | **p-values (wrt 0 min)** | **p-values (wrt 5 min)** |
| Normalized Area Fraction Rab 5 | 0 min | 15 | 122 | 1.00 | 0.00 | 0.00 | 1.00 | | 1.23E-16 |
| | 5 min | 15 | 122 | 1.42 | 0.92 | 0.08 | 1.26 | 1.23E-16 | |
| | 10 min | 15 | 122 | 1.40 | 0.59 | 0.05 | 1.31 | 2.81E-20 | 0.75 |
| | 15 min | 15 | 122 | 1.61 | 0.99 | 0.09 | 1.49 | 2.90E-23 | |

| FigS6d | | | | | | | | | | |
|---|---|---|---|---|---|---|---|---|---|---|
| **Parameters** | **Conditions** | **$N_{cell}$** | **$n_{ROI}$** | **Mean** | **SD** | **SEM** | **Median** | **p-values (wrt 0 min)** | **p-values (wrt 5 min)** | **p-values (wrt 10 min)** |
| Normalized Area Fraction Rab 4 | 0 min | 43 | 68 | 1.00 | 0.00 | 0.00 | 1.00 | | 1.00 | 2.63E-19 |

| | | | | | | | | | | |
|---|---|---|---|---|---|---|---|---|---|---|
| | 5 min | 43 | 68 | 1.05 | 0.35 | 0.04 | 1.00 | 1.00 | | 0.36 |
| | 10 min | 43 | 68 | 1.15 | 0.38 | 0.05 | 1.05 | 0.02 | 0.06 | |
| | 15 min | 43 | 68 | 1.13 | 0.47 | 0.06 | 1.02 | 0.21 | | 0.76 |

| **Fig S6e** | | | | | | | | | | |
|---|---|---|---|---|---|---|---|---|---|---|
| **Parameters** | **Conditions** | **$N_{cell}$** | **$n_{ROI}$** | **Mean** | **SD** | **SEM** | **Median** | **p-values (wrt 0 min)** | **p-values (wrt 3 min)** | **p-values (wrt 10 min)** |
| Area Fraction clathrin pits | 0 min | 74 | 74 | 0.64 | 0.42 | 0.05 | 0.59 | | 0.008 | 2.49E-03 |
| | 3 min | 82 | 82 | 0.77 | 0.37 | 0.04 | 0.79 | 0.008 | | 0.94 |
| | 6min | 94 | 94 | 0.73 | 0.42 | 0.04 | 0.72 | 0.14 | 0.33 | |

| **Fig S7b** | | | | | | | | | | |
|---|---|---|---|---|---|---|---|---|---|---|
| **Parameters** | **Conditions** | **$N_{cell}$** | **$n_{ROI}$** | **Mean** | **SD** | **SEM** | **Median** | **p-values (wrt 0 min)** | **p-values (wrt 3 min)** | **p-values (wrt 10 min)** |
| Area Fraction Rab 5 | 0 min (control) | 66 | 696 | 0.09 | 0.038 | 0.001 | 0.094 | | 7.00E-04 | |
| | 3 min(Control) | 58 | 562 | 0.101 | 0.043 | 0.002 | 0.099 | 7.00E-04 | | 2.81E-20 |
| | 6 min(Control) | 57 | 411 | 0.102 | 0.031 | 0.002 | 0.098 | 1.00E-04 | 0.62 | 0.75 |
| | 0 min(Dynasore) | 43 | 386 | 0.13 | 0.053 | 0.003 | 0.119 | | <1E-5 | |
| | 3 min(Dynasore) | 62 | 358 | 0.109 | 0.066 | 0.003 | 0.091 | <1E-5 | | 0.02 |
| | 6 min(Dynasore) | 50 | 187 | 0.106 | 0.059 | 0.004 | 0.096 | <1E-5 | 0.89 | |

| Parameters | Conditions | $N_{cell}$ | $n_{ROI}$ | Mean | SD | SEM | Median | p-values (wrt 0 min) | p-values (wrt 3 min) | p-values (wrt 10 min) |
|---|---|---|---|---|---|---|---|---|---|---|
| Area Fraction Rab 4 | 0 min (Control) | 125 | 125 | 0.339 | 0.025 | 0.002 | 0.337 | | 4.00E-04 | 0.02 |
| | 5 min (Control) | 129 | 129 | 0.352 | 0.03 | 0.003 | 0.352 | 4.00E-04 | | 0.06 |
| | 10 min (Control) | 70 | 70 | 0.301 | 0.058 | 0.007 | 0.304 | <1E-5 | <1E-5 | |
| | 0 min (Dynasore) | 206 | 206 | 0.27 | 0.06 | 0.004 | 0.27 | | <1E-5 | 0.49 |
| | 5 min (Dynasore) | 170 | 170 | 0.32 | 0.076 | 0.006 | 0.34 | <1E-5 | | |
| | 10 min (Dynasore) | 103 | 103 | 0.28 | 0.069 | 0.007 | 0.28 | 0.203 | 3.00E-05 | |

| **Fig S7d** | | | | | | | | | |
|---|---|---|---|---|---|---|---|---|---|
| **Parameters** | **Conditions** | $N_{cell}$ | $n_{FBR}$ | **Mean** | **SD** | **SEM** | **Median** | **p-values (wrt C)** | **p-values (wrt P2)** |
| $SD_{time}$ (nm) | C (Control) | 20 | 3184 | 6.88 | 2.43 | 0.04 | 6.59 | | <1E-5 |
| | P2 (Control) | 20 | 1859 | 8.47 | 2.56 | 0.06 | 8.44 | <1E-5 | |
| | P3 (Control) | 20 | 796 | 7.47 | 2.03 | 0.07 | 7.23 | <1E-5 | 0 |
| | C (Dynasore) | 11 | 8861 | 6.01 | 1.99 | 0.02 | | | <1E-5 |
| | P2 (Dynasore) | 11 | 4042 | 7.08 | 2.02 | 0.03 | 6.98 | <1E-5 | |
| | P3 (Dynasore) | 11 | 1733 | 5.42 | 1.57 | 0.04 | 5.17 | 0 | 0 |
| Tension (pN/µm) | C (Control) | 20 | 2006 | 106.1 | 266.6 | 5.95 | 40.7 | | 0 |
| | P2 (Control) | 20 | 957 | 33.05 | 35.41 | 1.14 | 22.79 | 0 | |
| | P3 (Control) | 20 | 355 | 45.8 | 82.57 | 4.38 | 27.76 | <1E-5 | <1E-5 |

| | | | | | | | | | |
|---|---|---|---|---|---|---|---|---|---|
| | C (Dynasore) | 11 | 6669 | 147.1 | 358.3 | 4.37 | 49.67 | | 0 |
| | P2 (Dynasore) | 11 | 3230 | 62.89 | 156.3 | 2.75 | 33.42 | 0 | |
| | P3 (Dynasore) | 11 | 1475 | 168.6 | 326.9 | 8.51 | 70.99 | <1E-5 | <1E-5 |
| $SD_{space}$ (nm) | C (Control) | 20 | 3184 | 9.17 | 3.63 | 0.06 | 8.84 | | <1E-5 |
| | P2 (Control) | 20 | 1859 | 9.62 | 3.27 | 0.08 | 9.37 | <1E-5 | |
| | P3 (Control) | 20 | 796 | 8.49 | 2.59 | 0.09 | 8.09 | 1.00E-04 | 0 |
| | C (Dynasore) | 11 | 8861 | 7.47 | 2.79 | 0.03 | 7.03 | | 0.02 |
| | P2 (Dynasore) | 11 | 4042 | 7.29 | 2.57 | 0.04 | 6.98 | 0.02 | |
| | P3 (Dynasore) | 11 | 1733 | 6.77 | 2.48 | 0.06 | 6.41 | 0 | <1E-5 |

| **Fig S7f** | | | | | | | | | |
|---|---|---|---|---|---|---|---|---|---|
| **Parameters** | **Conditions** | **$N_{cell}$** | **$n_{FBR}$** | **Mean** | **SD** | **SEM** | **Median** | **p-values (wrt C)** | **p-values (wrt P2)** |
| Normalized $SD_{space}$ | C (Control) | 20 | 20 | 1 | 0 | 0 | 1 | | 0.5728 |
| | P2 (Control) | 20 | 20 | 1.04 | 0.11 | 0.02 | 1.03 | 0.5728 | |
| | P3 (Control) | 20 | 20 | 0.95 | 0.15 | 0.03 | 0.98 | 0.5728 | 0.081 |
| | C (Dynasore) | 11 | 11 | 1 | 0 | 0 | 1 | | 6.00E-04 |
| | P2 (Dynasore) | 11 | 11 | 0.79 | 0.25 | 0.08 | 0.66 | 6.00E-04 | |
| | P3 (Dynasore) | 11 | 11 | 0.77 | 0.26 | 0.08 | 0.74 | 0.0076 | 0.8438 |

| **Fig S7g** | | | | | | | | |
|---|---|---|---|---|---|---|---|---|
| **Parameters** | **Conditions** | **$N_{cell}$** | **$n_{FBR}$** | **Mean** | **SD** | **SEM** | **Median** | **p-values (wrt Control)** |
| Activity | Control | 11 | 4302 | 4.00E-04 | 9.00E-04 | 1.00E-05 | 1.00E-04 | |

| | | | | | | | | |
|---|---|---|---|---|---|---|---|---|
| | Dynasore | 11 | 8861 | 3.00E-04 | 8.00E-04 | 9.00E-06 | 1.00E-04 | 0 |

| Fig S7h | | | | | | | | |
|---|---|---|---|---|---|---|---|---|
| Normalized Excess Area | Control | 6 | 3238 | 1.05 | 0.29 | 0.005 | 1 | |
| | ML 141 | 17 | 11397 | 1.1 | 0.36 | 0.003 | 1.03 | 7.79E-05 |

| Fig S7i | | | | | | | | | |
|---|---|---|---|---|---|---|---|---|---|
| **Parameters** | **Conditions** | **$N_{cell}$** | **$n_{FBR}$** | **Mean** | **SD** | **SEM** | **Median** | **p-values (wrt C)** | **p-values (wrt P2)** |
| $SD_{time}$ (nm) Cell Wise | C (Control) | 11 | | 6.94 | 2.01 | 0.61 | 6.51 | | 0.168 |
| | P2 (Control) | 11 | | 8.2 | 2.14 | 0.64 | 8.44 | 0.17 | |
| | P3 (Control) | 11 | | 7.3 | 1.49 | 0.45 | 7.09 | 0.47 | 0.24 |
| | C (Dynasore) | 11 | | 5.78 | 1.002 | 0.3 | 5.47 | | 0.02 |
| | P2 (Dyna) | 11 | | 6.89 | 0.83 | 0.25 | 7.09 | 0.02 | |
| | P3 (Dyna) | 11 | | 5.4 | 0.77 | 0.23 | 5.62 | 0.55 | 0.001 |
| $SD_{space}$ (nm) Cell Wise | C (Control) | 11 | | 9.54 | 2.75 | 0.83 | 8.78 | | 0.84 |
| | P2 (Control) | 11 | | 9.48 | 2.501 | 0.75 | 8.06 | 0.84 | |
| | P3 (Control) | 11 | | 8.43 | 1.77 | 0.54 | 8.13 | 0.39 | 0.26 |
| | C (Dynasore) | 11 | | 7.035 | 1.13 | 0.34 | 6.99 | | 0.95 |
| | P2 (Dyna) | 11 | | 7.039 | 0.88 | 0.27 | 7.18 | 0.95 | |
| | P3 (Dyna) | 11 | | 6.83 | 1.32 | 0.4 | 7.35 | 0.79 | 1 |
| Tension (pN/µm) Cell Wise | C (Control) | 11 | | 44.32 | 22.28 | 6.72 | 43.7 | | 0.02 |
| | P2 (Control) | 11 | | 30.93 | 24.55 | 7.4 | 23.14 | 0.02 | |
| | P3 (Control) | 11 | | 32.78 | 14.75 | 4.45 | 32.5 | 0.26 | 0.36 |
| | C (Dynasore) | 11 | | 55.54 | 21.65 | 6.53 | 56.05 | | 0.015 |
| | P2 (Dyna) | 11 | | 35.96 | 14.14 | 4.26 | 32.6 | 0.015 | |

| | | | | | | | | | |
|---|---|---|---|---|---|---|---|---|---|
| | P3 (Dyna) | 11 | | 74.05 | 33.05 | 9.96 | 63.2 | 0.17 | 5.00E-04 |

| Fig S8b | | | | | | | | | |
|---|---|---|---|---|---|---|---|---|---|
| **Parameters** | **Conditions** | **$N_{cell}$** | **$n_{FBR}$** | **Mean** | **SD** | **SEM** | **Median** | **p-values (wrt Control)** | **p-values (wrt P2)** |
| $SD_{time}$ (nm) | C (Control) | 9 | 8188 | 3.64 | 0.98 | 0.01 | 3.47 | | 4.24E-73 |
| | P2 (Control) | 9 | 3941 | 4.03 | 1.17 | 0.02 | 3.82 | 4.24E-73 | |
| | P3 (Control) | 9 | 1145 | 3.59 | 1.03 | 0.03 | 3.44 | 0.04 | 0 |
| | C (Dyna KO) | 15 | 12524 | 3.48 | 1.11 | 0.01 | 3.24 | | 3.57E-134 |
| | P2 (Dyna KO) | 15 | 5476 | 3.91 | 1.26 | 0.01 | 3.72 | 3.57E-134 | |
| | P3 (Dyna KO) | 15 | 2033 | 3.35 | 1.01 | 0.02 | 3.19 | 3.76E-5 | 0 |
| **Parameters** | **Conditions** | **$N_{cell}$** | **$n_{FBR}$** | **Mean** | **SD** | **SEM** | **Median** | **p-values (wrt Control)** | **p-values (wrt P2)** |
| Tension (pN/μm) | C (Control) | 9 | 5368 | 410.38 | 677.40 | 9.25 | 161.37 | | 0 |
| | P2 (Control) | 9 | 2677 | 281.64 | 523.65 | 10.12 | 118.33 | 0 | |
| | P3 (Control) | 9 | 711 | 380.5 | 693.63 | 26.01 | 135.39 | 1.32E-11 | 5.62E-43 |
| | C (Dyna KO) | 15 | 8192 | 444.64 | 696.32 | 7.69 | 188.99 | | 0 |
| | P2 (Dyna KO) | 15 | 3494 | 370.16 | 674.46 | 11.41 | 130.48 | 0 | |
| | P3 (Dyna KO) | 15 | 1331 | 570.48 | 879.23 | 24.1 | 240.87 | 7.67108E-5 | 5.23E-5 |

| Fig S9b | | | | | | | | | |
|---|---|---|---|---|---|---|---|---|---|
| **Parameters** | **Conditions** | **$N_{cell}$** | **$n_{patches}$** | **Mean** | **SD** | **SEM** | **Median** | **p-value (wrt 0 min )** | **p-value (wrt 3 min)** |
| Area Fraction Rab 5(ATP Dep) | 0 min | 26 | 168 | 0.09 | 0.06 | 0.005 | 0.08 | | 0.07 |
| | 3 min | 41 | 288 | 0.09 | 0.05 | 0.003 | 0.09 | 0.07 | |

| | | | | | | | | | |
|---|---|---|---|---|---|---|---|---|---|
| | 6min | 39 | 255 | 0.08 | 0.05 | 0.003 | 0.09 | 0.52 | 0.17 |

| **Fig S9c** | | | | | | | | | |
|---|---|---|---|---|---|---|---|---|---|
| **Parameters** | **Conditions** | **$N_{cell}$** | | **Mean** | **SD** | **SEM** | **Median** | **p-values (wrt C)** | **p-values (wrt P2)** |
| $SD_{time}$ (nm) Cell wise | C (ATP Dep.) | 9 | | 3.199 | 0.57 | 0.19 | 3.35 | | 6.00E-04 |
| | P2 (ATP Dep.) | 9 | | 6.247 | 1.34 | 0.45 | 6.22 | 6.00E-04 | |
| | P3 (ATP Dep.) | 9 | | 3.499 | 0.65 | 0.22 | 3.23 | 0.54 | 6.00E-04 |
| | C (Chol. Dep.) | 16 | | 5.598 | 0.84 | 0.211 | 5.39 | | 1.00E-05 |
| | P2(Chol.Dep.) | 16 | | 7.49 | 0.69 | 0.17 | 7.56 | 1.00E-05 | |
| | P3(Chol.Dep.) | 16 | | 6.3 | 0.785 | 0.196 | 6.44 | 0.027 | 1.00E-04 |
| | C (ATP Dep. +Chol. Dep.) | 15 | | 4.74 | 0.63 | 0.16 | 4.73 | | $<10^{-5}$ |
| | P2 (ATP Dep. +Chol. Dep.) | 15 | | 7.628 | 0.9 | 0.23 | 7.5 | $<10^{-5}$ | |
| | P3 (ATP Dep. +Chol. Dep.) | 15 | | 8.014 | 2.14 | 0.55 | 8.44 | 9.66E-05 | 0.36 |
| **Parameters** | **Conditions** | **$N_{cell}$** | | **Mean** | **SD** | **SEM** | **Median** | **p-values (wrt C)** | **p-values (wrt P2)** |
| $SD_{space}$ (nm) Cell wise | C (ATP Dep.) | 9 | | 9.79 | 0.8 | 0.27 | 9.63 | | 0.02 |
| | P2 (ATP Dep.) | 9 | | 8.14 | 1.26 | 0.42 | 8.26 | 0.02 | |
| | P3 (ATP Dep.) | 9 | | 7.13 | 0.71 | 0.24 | 7.54 | 0 | 0.09 |
| | C (Chol. Dep.) | 16 | | 6.72 | 0.67 | 0.17 | 6.66 | | 0.78 |
| | P2(Chol.Dep.) | 16 | | 6.83 | 1.19 | 0.3 | 6.98 | 0.78 | |
| | P3(Chol.Dep.) | 16 | | 6.75 | 0.71 | 0.18 | 6.87 | 0.4 | 0.87 |
| | C (ATP Dep. +Chol. Dep.) | 15 | | 7.83 | 0.44 | 0.11 | 7.8 | | 0.17 |
| | P2 (ATP Dep. +Chol. Dep.) | 15 | | 7.41 | 0.96 | 0.25 | 7.35 | 0.17 | |
| | P3 (ATP Dep. +Chol. Dep.) | 15 | | 7.95 | 1.95 | 0.5 | 8.13 | 0.36 | 0.2 |
| Tension (pN/µm) Cell Wise | C (ATP Dep.) | 9 | | 270.73 | 236.32 | 78.77 | 163.86 | | 6.00E-04 |
| | P2 (ATP Dep.) | 9 | | 61.27 | 33.21 | 11.07 | 46.28 | 6.00E-04 | |
| | P3 (ATP Dep.) | 9 | | 163.87 | 58.797 | 19.599 | 175.56 | 0.479 | 0.001 |
| | C (Chol. Dep.) | 16 | | 54.7 | 16.997 | 4.25 | 55.12 | | 1.00E-04 |

| | | | | | | | | | |
|---|---|---|---|---|---|---|---|---|---|
| | P2(Chol.Dep.) | 16 | | 31.87 | 6.19 | 1.55 | 32.04 | 1.00E-04 | |
| | P3 Chol.Dep.) | 16 | | 43.84 | 13.32 | 3.33 | 40.12 | 0.09 | 0.002 |
| | C (ATP Dep. +Chol. Dep.) | 15 | | 110.48 | 70.796 | 18.28 | 95.78 | | $<10^{-5}$ |
| | P2 (ATP Dep. +Chol. Dep.) | 15 | | 37.497 | 12.24 | 3.16 | 32.6 | $<10^{-5}$ | |
| | P3 (ATP Dep. +Chol. Dep.) | 15 | | 42.13 | 27.16 | 7.01 | 32.94 | 1.00E-04 | 0.56 |

| Fig S9d | | | | | | | | | |
|---|---|---|---|---|---|---|---|---|---|
| **Parameters** | **Conditions** | **$N_{cell}$** | **$n_{FBR}$** | **Mean** | **SD** | **SEM** | **Median** | **p-values (wrt C)** | **p-values (wrt P2)** |
| $SD_{time}$ (nm) | C (ATP Dep.) | 9 | 1962 | 3.09 | 0.85 | 0.02 | 2.92 | | 0 |
| | P2 (ATP Dep.) | 9 | 1271 | 6.31 | 1.63 | 0.05 | 6.22 | 0 | |
| | P3 (ATP Dep.) | 9 | 550 | 3.33 | 0.81 | 0.03 | 3.19 | <1E-5 | 0 |
| | C (Chol. Dep.) | 16 | 8072 | 5.41 | 1.5 | 0.02 | 5.21 | | 0 |
| | P2 (Chol. Dep.) | 16 | 3438 | 7.62 | 1.83 | 0.03 | 7.47 | 0 | |
| | P3 (Chol. Dep.) | 16 | 1366 | 6.61 | 1.51 | 0.04 | 6.43 | <1E-5 | 0 |
| | C (ATP Dep. +Chol. Dep.) | 15 | 8351 | 4.86 | 1.34 | 0.01 | 4.68 | | 0 |
| | P2 (ATP Dep. +Chol. Dep.) | 15 | 4681 | 7.74 | 2.12 | 0.03 | 7.51 | 0 | |
| | P3 (ATP Dep. +Chol. Dep.) | 15 | 1966 | 7.7 | 2.52 | 0.06 | 7.36 | 0 | 0.05 |
| $SD_{space}$ (nm) | C (ATP Dep.) | 9 | 1962 | 9.88 | 2.11 | 0.05 | 9.8 | | 0 |
| | P2 (ATP Dep.) | 9 | 1271 | 8.26 | 2.07 | 0.06 | 8.07 | 0 | |
| | P3 (ATP Dep.) | 9 | 550 | 7.46 | 1.65 | 0.07 | 7.42 | 0 | <1E-5 |
| | C (Chol. Dep.) | 16 | 8072 | 6.89 | 2.34 | 0.03 | 6.44 | | 0.15 |
| | P2 (Chol. Dep.) | 16 | 3438 | 6.99 | 2.51 | 0.04 | 6.57 | 0.15 | |
| | P3 (Chol. Dep.) | 16 | 1366 | 7.07 | 2.19 | 0.06 | 6.71 | 3.84E-05 | 0.01 |
| | C (ATP Dep. +Chol. Dep.) | 15 | 8351 | 8.25 | 2.8 | 0.03 | 7.81 | | <1E-5 |
| | P2 (ATP Dep. +Chol. Dep.) | 15 | 4681 | 8.02 | 2.83 | 0.04 | 7.54 | <1E-5 | |

| | | | | | | | | | |
|---|---|---|---|---|---|---|---|---|---|
| | P3 (ATP Dep. +Chol. Dep.) | 15 | 1966 | 8.04 | 2.72 | 0.06 | 7.59 | 0.004 | 0.45 |
| **Parameters** | **Conditions** | **$N_{cell}$** | **$n_{FBR}$** | **Mean** | **SD** | **SEM** | **Median** | **p-values (wrt C)** | **p-values (wrt P2)** |
| Relative Tension | C (ATP Dep.) | 9 | 1363 | 2.77 | 4.65 | 0.13 | 1 | | 0 |
| | P2 (ATP Dep.) | 9 | 1014 | 0.37 | 0.48 | 0.02 | 0.24 | 0 | |
| | P3 (ATP Dep.) | 9 | 390 | 1.31 | 1.97 | 0.1 | 0.75 | <1E-5 | <1E-5 |
| | C (Chol. Dep.) | 16 | 6032 | 2.32 | 6.01 | 0.08 | 1 | | 0 |
| | P2 (Chol. Dep.) | 16 | 2291 | 0.71 | 0.83 | 0.02 | 0.53 | 0 | |
| | P3 (Chol. Dep.) | 16 | 1087 | 1 | 1.69 | 0.05 | 0.69 | 0 | <1E-5 |
| | C (ATP Dep. +Chol. Dep.) | 15 | 7169 | 2.68 | 5.07 | 0.06 | 1 | | 0 |
| | P2 (ATP Dep. +Chol. Dep.) | 15 | 4281 | 0.59 | 0.91 | 0.01 | 0.4 | 0 | |
| | P3 (ATP Dep. +Chol. Dep.) | 15 | 1677 | 0.97 | 2.48 | 0.06 | 0.41 | 0 | 0.002 |

| **Table S2** | | | |
|---|---|---|---|
| Values obtained from LMM analysis of FBR-wise comparisons | | | |
| **Normal Slow - tension** | | | |
| | **p-value** | **slope** | **Standard error** |
| C-P2 | 0.007 | -0.26 | 0.1 |
| C-P3 | $<10^{-5}$ | -0.55 | 0.1 |
| P2-P3 | 0.001 | -0.29 | 0.09 |
| **Normal Slow - $SD_{time}$** | | | |
| C-P2 | 0.009 | 0.07 | 0.03 |
| C-P3 | $<10^{-5}$ | 0.16 | 0.03 |
| P2-P3 | 0.001 | 0.09 | 0.03 |
| **Normal Slow - $SD_{space}$** | | | |
| C-P2 | 0.262 | 0.02 | 0.02 |
| C-P3 | 0.132 | 0.04 | 0.03 |
| P2-P3 | 0.509 | 0.02 | 0.03 |
| | **p-value** | **slope** | **Standard error** |

| | | | |
|---|---|---|---|
| **Normal Fast - Tension** | | | |
| C-P2 | 0.001 | -0.44 | 0.14 |
| C-P3 | 0.0003 | -0.35 | 0.1 |
| P2-P3 | 0.492 | 0.1 | 0.14 |
| **Normal Fast - $SD_{time}$** | | | |
| C-P2 | 0.0001 | 0.15 | 0.04 |
| C-P3 | 0.0009 | 0.13 | 0.04 |
| P2-P3 | 0.672 | -0.02 | 0.04 |
| **Normal Fast - $SD_{space}$** | | | |
| C-P2 | 0.921 | 0 | 0.04 |
| C-P3 | 0.785 | -0.01 | 0.02 |
| P2-P3 | 0.562 | -0.02 | 0.03 |
| **Dynasore – tension** | | | |
| C-P2 | 0.002 | -0.38 | 0.13 |
| C-P3 | 0.019 | 0.32 | 0.14 |
| P2-P3 | $<10^{-5}$ | 0.7 | 0.11 |
| **Dynasore - $SD_{time}$** | | | |
| C-P2 | 0.0005 | 0.14 | 0.04 |
| C-P3 | 0.109 | -0.07 | 0.05 |
| P2-P3 | 2.28E-08 | -0.21 | 0.04 |
| **Dynasore - $SD_{space}$** | | | |
| C-P2 | 0.648 | 0.01 | 0.03 |
| C-P3 | 0.324 | -0.04 | 0.04 |
| P2-P3 | 0.112 | -0.06 | 0.03 |
| **Cyto D - $SD_{time}$** | | | |
| C-P2 | 0.007 | 0.06 | 0.02 |
| C-P3 | 0.642 | 0.01 | 0.02 |
| P2-P3 | 0.014 | -0.05 | 0.02 |
| **Cyto D - $SD_{space}$** | | | |
| C-P2 | 0.13 | -0.03 | 0.02 |
| C-P3 | 0.005 | -0.07 | 0.03 |
| P2-P3 | 0.031 | -0.04 | 0.02 |
| | **p-value** | **slope** | **Standard error** |
| **Cyto D - tension** | | | |
| C-P2 | 0.576 | -0.04 | 0.07 |
| C-P3 | 0.965 | 0 | 0.06 |

| | | | |
|---|---|---|---|
| P2-P3 | 0.562 | 0.04 | 0.07 |
| **ATP depletion - $SD_{time}$** | | | |
| C-P2 | 3.74E-10 | 0.54 | 0.09 |
| C-P3 | 0.0005 | 0.19 | 0.05 |
| P2-P3 | 0.0001 | -0.35 | 0.09 |
| **ATP depletion - $SD_{space}$** | | | |
| C-P2 | 1.46E-12 | -0.17 | 0.02 |
| C-P3 | 8.16E-20 | -0.35 | 0.04 |
| P2-P3 | 0.0018 | -0.18 | 0.06 |
| **ATP depletion - tension** | | | |
| C-P2 | 1.49E-31 | -1.55 | 0.13 |
| C-P3 | 0.0003 | -0.62 | 0.17 |
| P2-P3 | 1.81E-07 | 0.93 | 0.18 |
| **ATP+ cholesterol depletion - $SD_{time}$** | | | |
| C-P2 | 1.61E-40 | 0.47 | 0.04 |
| C-P3 | 1.16E-33 | 0.54 | 0.04 |
| P2-P3 | 0.123 | 0.07 | 0.05 |
| **ATP+ cholesterol depletion - $SD_{space}$** | | | |
| C-P2 | 3.75E-08 | -0.73 | 0.13 |
| C-P3 | 3.45E-09 | -0.9 | 0.15 |
| P2-P3 | 0.0002 | -0.17 | 0.05 |
| **ATP+ cholesterol depletion – tension** | | | |
| C-P2 | 7.58E-44 | -1.16 | 0.08 |
| C-P3 | 1.74E-62 | -1.3 | 0.08 |
| P2-P3 | 0.024 | -0.15 | 0.07 |
| **Cholesterol Depletion - tension** | | | |
| C-P2 | 1.17E-18 | -0.57 | 0.06 |
| C-P3 | 0.0003 | -0.25 | 0.07 |
| P2-P3 | 1.07E-10 | 0.31 | 0.05 |
| | **p-value** | **slope** | **Standard error** |
| **Cholesterol Depletion - $SD_{time}$** | | | |
| C-P2 | 1.30E-22 | 0.28 | 0.03 |
| C-P3 | 0.0003 | 0.12 | 0.03 |
| P2-P3 | 6.79E-10 | -0.16 | 0.03 |

| | | | |
|---|---|---|---|
| **Cholesterol Depletion - $SD_{space}$** | | | |
| C-P2 | 0.279 | 0.03 | 0.03 |
| C-P3 | 0.742 | -0.01 | 0.02 |
| P2-P3 | 0.235 | -0.03 | 0.03 |
| **Dynamin Mutant $SD_{time}$** | | | |
| C-P2 | 2.25E-05 | 0.13 | 0.03 |
| C-P3 | 0.46 | -0.03 | 0.04 |
| P2-P3 | 2.82E-05 | -0.16 | 0.04 |
| **Dynamin Mutant Tension** | | | |
| C-P2 | 5.19E-03 | -0.29 | 0.10 |
| C-P3 | 0.03 | 0.24 | 0.11 |
| P2-P3 | 4.17E-05 | 0.53 | 0.13 |
| **Excess Area** | | | |
| **C(Ctrl)-C(Dynasore)** | | | |
| 7 sets of ctrl, 3 sets of dynasore | 6.43E-11 | 0.22 | 0.03 |

### Extended Methods:

#### Details about Linear Mixed Model:

We implement a Linear Mixed Model (LMM) using the following formula (Eq. 1):

Value ~ Phase +(Phase|Set:Cell)' (Eq 1)

The measured parameter (Value, representing $SD_{time}$ or $SD_{space}$ log (excess area) or log(tension)) was considered as a function of de-adhesion phase. Predictor variables - set number of experiment (Set) and cell number (Cell) - are grouping variables. Cell nested in Set contribute random intercepts (with possible correlation with random slope) to the model to control for the variation across sets, treat cells to be grouped under sets and FBRs to be grouped under cells and thus account of the repeated measurements in clusters. The Linear Mixed Model class from the Statistics Toolbox in MATLAB was used for model fitting. The linear mixed model coefficients were estimated using maximal likelihood (ML) as the default settings.
